# DARPP-32 and t-DARPP promote lung cancer growth through IKKα-dependent cell migration and Akt/Erk-mediated cell survival

**DOI:** 10.1101/229658

**Authors:** Sk. Kayum Alam, Matteo Astone, Ping Liu, Stephanie R. Hall, Abbygail M. Coyle, Erin N. Dankert, Dane K. Hoffman, Wei Zhang, Rui Kuang, Anja C. Roden, Aaron S. Mansfield, Luke H. Hoeppner

**Author notes:** These second authors contributed equally (Matteo Astone, Ping Liu). Corresponding Author: Luke H. Hoeppner, Ph.D. The Hormel Institute, University of Minnesota, 801 16th Avenue NE, Austin, MN 55912.

## Abstract

Lung cancer is the leading cause of cancer-related death worldwide. In this study, we demonstrate that elevated expression of dopamine and cyclic adenosine monophosphate-regulated phosphoprotein, Mr 32000 (DARPP-32) and its truncated splice variant t-DARPP promotes lung tumor growth, while abrogation of DARPP-32 expression in human non-small cell lung cancer (NSCLC) cells reduces tumor growth in orthotopic mouse models. We observe a novel physical interaction between DARPP-32 and inhibitory kappa B kinase-α (IKKα) that promotes NSCLC cell migration through non-canonical nuclear factor kappa-light-chain-enhancer of activated B cells 2 (NF-κB2) signaling. Bioinformatics analysis of 513 lung adenocarcinoma patients reveals elevated t-DARPP isoform expression is associated with poor overall survival. Histopathological investigation of 62 human lung adenocarcinoma tissues also showed that t-DARPP expression is elevated with increasing tumor (T) stage. Our data suggest that DARPP-32 is a negative prognostic marker associated with increasing stages of NSCLC and may represent a novel therapeutic target.

## Introduction

Lung cancer is the leading cause of cancer deaths among both men and women (Torre et al., 2016). In 2017, an estimated 160,420 lung cancer deaths will occur in the United States (Siegel et al., 2017). Nonsmall cell lung cancer (NSCLC) represents 85-90% of all cases of lung cancer and carries a very poor survival rate with less than 15% of patients surviving more than five years (Cetin et al., 2011; Molina et al., 2008). Despite administration of standard chemotherapeutic agents with evolving systemic cancer therapies directed at driver mutations (EGFR, BRAF and ALK), inhibiting angiogenesis (anti-VEGF therapy) and immune-checkpoint blockade (anti-PD-1 antibody), these statistics remain dismal due to the large number of patients diagnosed with advanced stage disease and the primary and secondary resistance to current therapies. A better understanding of the mechanisms that regulate lung tumor growth, metastasis and drug resistance will result in new diagnostic tools and therapeutic strategies to improve the clinical outlook and quality of life of patients afflicted with this deadly disease.

Dopamine and cyclic adenosine monophosphate-regulated phosphoprotein, Mr 32000 (DARPP-32) is an effector molecule that plays an important role in dopaminergic neurotransmission. This 32-kDa protein was initially discovered in the neostriatum in the brain as substrate of dopamine-activated protein kinase A (PKA) (Walaas et al., 1983). Phosphorylation at threonine-34 (T34) by PKA causes DARPP-32-mediated inhibition of protein phosphatase-1 (PP-1) (Hemmings et al., 1984), hence DARPP-32 is also called phosphoprotein phosphatase-1 regulatory subunit 1B (PPP1R1B). DARPP-32 is converted to an inhibitor of PKA upon phosphorylation of its T75 residue by cyclin-dependent kinase 5 (Cdk5) (Bibb et al., 1999). The ability of DARPP-32 to function as either a kinase or a phosphatase inhibitor enables it to precisely modulate dopaminergic neurotransmission (Bibb et al., 1999; Greengard, 2001).

In the early 2000s, El-Rifai and colleagues discovered DARPP-32 is frequently amplified and upregulated in gastric cancer (Belkhiri et al., 2005; El-Rifai et al., 2002). Cloning and sequence assembly analysis revealed a novel transcriptional splice variant of DARPP-32 is also overexpressed in gastric cancer. The N-terminally truncated isoform of DARPP-32, termed t-DARPP, was found to utilize a unique alternative first exon located within intron 1 of DARPP-32 and to lack the first 36 amino acids of DARPP-32, including the T34 phosphorylation residue required for DARPP-32-mediated PP-1 inhibition (El-Rifai et al., 2002). Overexpression of both DARPP-32 and t-DARPP has been observed in 68% of gastric cancers (Belkhiri et al., 2005; El-Rifai et al., 2002). Elevated expression levels of DARPP-32 and t-DARPP have also been associated with many adenocarcinomas, including stomach, colon, prostate and breast cancers (Beckler et al., 2003; Belkhiri et al., 2012; Christenson et al., 2015; Gu et al., 2009; Vangamudi et al., 2010; Wang et al., 2005). Reports have implicated DARPP-32 and t-DARPP in cancer cell proliferation, survival, invasion and angiogenesis (Belkhiri et al., 2016). Several studies have demonstrated that DARPP-32 and t-DARPP protect cancer cells from drug-induced apoptosis, which is dependent upon their T75 phosphorylation residue (Belkhiri et al., 2005; El-Rifai et al., 2002) and involves upregulation of Akt and Bcl2 proteins (Belkhiri, Dar, Zaika, et al., 2008; Belkhiri et al., 2012; Zhu et al., 2011). To date, the role of DARPP-32 isoforms in lung cancer remains unexplored. However, we recently described the role of dopamine signaling in NSCLC by demonstrating dopamine D2 receptor agonists inhibit lung cancer growth by reducing angiogenesis and tumor infiltrating myeloid derived suppressor cells in preclinical orthotopic murine models (Hoeppner et al., 2015). Given the role of dopamine signaling in lung cancer and the oncogenic nature of DARPP-32 isoforms in a variety of tumor types, we sought to determine whether DARPP-32 and t-DARPP contribute to lung cancer growth, progression and drug resistance.

Nuclear factor kappa-light-chain-enhancer of activated B cells (NF-κB) is a transcription factor that regulates numerous biological processes, such as immunity, inflammation, cell growth, differentiation, migration, tumorigenesis and apoptosis (Hayden et al., 2008). The family of NF-κB proteins is comprised of structurally homologous transcription factors, including NF-κB1 (p105/50), NF-κB2 (p100/52), RelA (p65), RelB and c-Rel (Caamano et al., 2002). In the absence of external stimuli, NF-κB proteins are sequestered in the cytoplasm by specific inhibitory proteins, inhibitors of NF-κB (IκBs) (Hayden et al., 2008). When a cell receives appropriate stimuli, IκB kinase (IKK) phosphorylation is initiated, leading to proteasome-mediated processing of p105 and p100. This cleavage event generates their respective mature proteins, p50 and p52, resulting in the nuclear translocation of previously sequestered NF-κB members (Beinke et al., 2004). NF-κB signaling has been categorized into canonical and non-canonical (i.e. alternative) pathways. Recent studies have shown that both canonical and non-canonical NF-κB pathways are capable of promoting oncogenesis by interacting with other cellular pathways in breast cancer, pancreatic ductal adenocarcinoma (PDAC) and glioblastomas (Bang et al., 2013; Kendellen et al., 2014; Rinkenbaugh et al., 2016). NF-κB1 pathway activation causes induction of the IKK complex that contains two catalytic subunits, IKKα and IKKβ and one scaffold subunit called nuclear factor κB essential modulator (NEMO) or IKKγ (Z. J. Chen et al., 1996; DiDonato et al., 1997). Dysregulation of the IKK complex can initiate constitutive activation of the NF-κB1 pathway in cancer cells (Baldwin, 2001). Non-canonical NF-κB2 signaling requires IKKα to mediate p100 cleavage into p52, but does not depend upon IKKβ and NF-κB essential modulator (NEMO), which are essential for canonical NF-κB1 signal transduction (Dejardin et al., 2002; Senftleben et al., 2001). A recent finding has suggested that constitutive activation of KRAS and IKK/NF-κB1 pathways expedites tumorigenesis and worsens survival in PDAC patients (Ling et al., 2012). Ablation of constitutive IKK activity by small molecule inhibitor reduces cellular NF-κB1 activity and melanoma cell survival *in vitro* and *in vivo* (Yang et al., 2006). A recent report has suggested that proinflammatory *H. pylori* infection and canonical NF-κB 1 activation play a significant role in the regulation DARPP-32 expression, which has been shown to counteract infection-induced cell death and promote cell survival in gastric carcinogenesis (Zhu, Soutto, et al., 2016).

We aimed to investigate the role of DARPP-32 isoforms in NSCLC. Here we demonstrate that DARPP-32 and t-DARPP promote cell survival and non-canonical NF-κB2 p52-mediated cell migration in lung cancer. In NSCLC patients, elevated expression of t-DARPP was found to be associated with tumor stage and worsened patient survival.

## Results

### DARPP-32 and t-DARPP promote cell survival in NSCLC through activation of Akt and Erk signaling

Given the oncogenic role of DARPP-32 in gastric and breast cancer progression (Belkhiri et al., 2005; Christenson et al., 2014; Vangamudi et al., 2010), we sought to determine whether DARPP-32 proteins regulate cell survival in NSCLC. First, we stably silenced endogenous DARPP-32 protein expression through lentiviral shRNA-mediated knockdown in A549 and H1650 human lung adenocarcinoma cells as well as H226 human lung squamous cell carcinoma cells (Fig. 1a, b, c). Two shRNAs targeting distinct regions of DARPP-32 were utilized to decrease the likelihood of potentially confounding off-target effects (Fig. 1a, b, c). To determine the role of DARPP-32 in regulation of cell survival, we first assessed apoptosis through flow cytometry-based annexin V assays. We observed increased annexin V positive cells in DARPP-32 knockdown cell lines compared to controls (Fig. 1d, e, f) suggesting that DARPP-32 inhibits apoptosis in lung cancer cells. Based on this finding, we next performed a colorimetric cell viability assay in A549 and H226 cells stably transduced with retrovirus to overexpress exogenous DARPP-32 proteins (Fig. 1g, h). Cell viability was significantly increased in DARPP-32 overexpressing cells compared to controls (Fig. 1i, j). An N-terminally truncated isoform and transcriptional variant of DARPP-32, called t-DARPP, lacks the protein phosphate inhibitory (PP-1) domain, which is phosphorylated at threonine 34 (T34) and important for dopamine signaling function (El-Rifai et al., 2002). Overexpression of t-DARPP in A549 and H226 lung cancer cells increased viability (Fig. 1i, j), suggesting that the N-terminal T34-dependent PP-1 regulatory function of DARPP-32 (Huang et al., 1999) does not contribute to regulation of cell viability. Given the role of t-DARPP in promoting cellular proliferation in gastrointestinal cancer (Vangamudi et al., 2011), we sought to determine whether DARPP-32 and t-DARPP proteins regulate proliferation of NSCLC cells. We found modulation of DARPP-32 isoforms does not alter proliferation of lung cancer cells using flow cytometry-based BrdU cell proliferation assays upon silencing endogenous DARPP-32 and overexpression of DARPP-32 and t-DARPP (Supplementary Fig. 1). Taken together, our findings suggest that DARPP-32 and t-DARPP promote lung tumor cell survival by regulating apoptosis but do not control cellular proliferation.

**Fig. 1:**
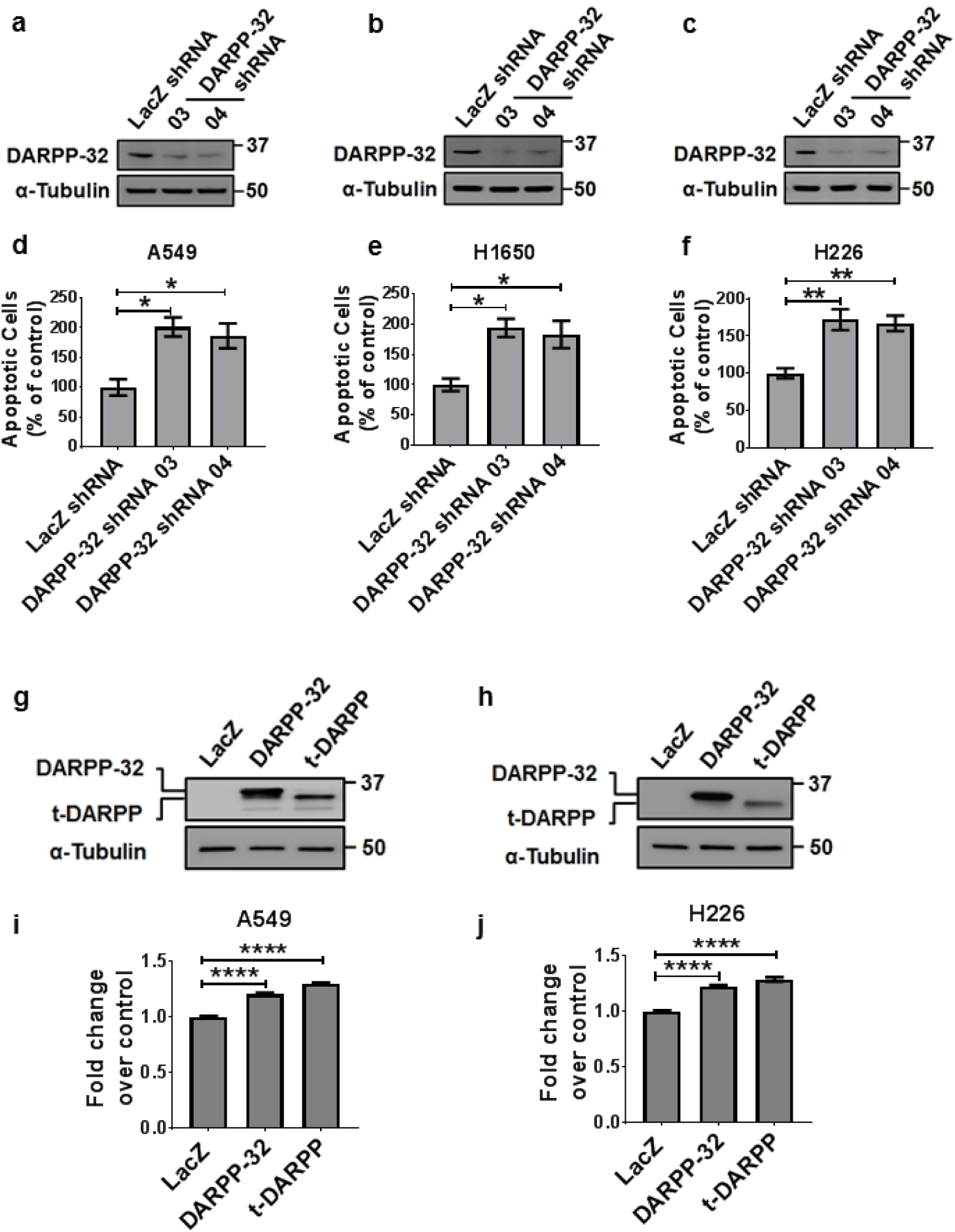
DARPP-32 promotes cell survival and negatively regulates apoptosis. **a** NSCLC A549, **b** H1650 and **c** H226 cell lines were transduced with lentivirus encoding LacZ control shRNA or DARPP-32 shRNAs (clone numbers 03 and 04). DARPP-32 and α-tubulin (loading control) proteins were detected by immunoblotting of cell lysates. **d** A549, **e** H1650 and **f** H226 cells transduced with control or DARPP-32 shRNAs were seeded into 60-mm culture dishes for 16h. Flow cytometry-based apoptosis assays were performed following incubation with anti-annexin V antibodies conjugated with APC. **g** A549 and h H226 cells were transduced with retrovirus containing control (LacZ), DARPP-32 or t-DARPP overexpressing clones. DARPP-32, t-DARPP and α-tubulin (loading control) proteins were detected by immunoblotting of cell lysates. **i** A549 and **j** H226 cells transduced with control, DARPP-32 or t-DARPP overexpressing clones were seeded into 96-well cell culture plates for 72h. Colorimeter-based cell survival assay was conducted using MTS reagents. Error bars indicate standard error of mean (SE). \**P*<0.05, \*\**P*<0.01 and \*\*\*\**P*<0.0001, one-way ANOVA.

To elucidate the molecular mechanism through which DARPP-32 proteins control NSCLC cell survival, we investigated Akt and Erk1/2 signaling as both pathways have been previously implicated in DARPP-32-mediated regulation of cell survival (Belkhiri, Dar, Zaika, et al., 2008; Hoeppner et al., 2015; Vangamudi et al., 2010). Ablation of DARPP-32 decreased phosphorylated Akt (p-Akt) and phosphorylated Erk (p-Erk) levels substantially, while corresponding total Akt and Erk1/2 protein expression remained unchanged by immunoblotting (Fig. 2a, b). Correspondingly, DARPP-32 overexpression resulted in increased phosphorylation of Akt and Erk1/2 (Fig. 2c, d). We found exogenous overexpression of t-DARPP and mutant DARPP-32 (T34A) proteins also elevates p-Akt and p-Erk1/2 levels, suggesting PP-1 activation by DARPP-32 T34 phosphorylation is not directly involved in stimulation of Akt and Erk signaling (Fig. 2c, d). Collectively, our results suggest that DARPP-32 promotes cell survival in a PP-1 independent manner through Akt and Erk1/2 signaling in NSCLC cells.

**Fig. 2:**
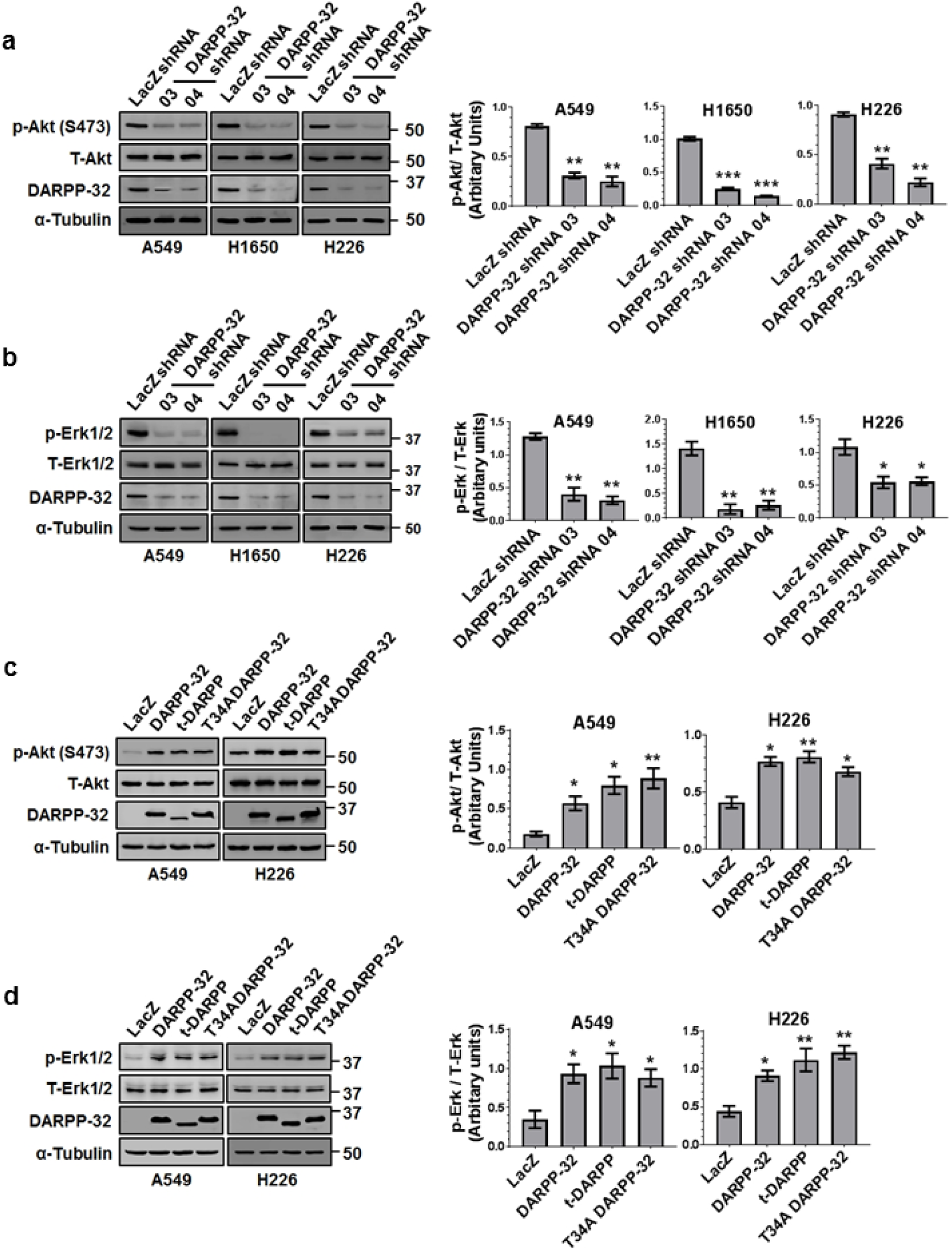
DARPP-32 regulates cell survival through Akt and Erk1/2. **a** A549, H1650 and H226 cells were transduced with control or DARPP-32 shRNAs. Cell lysates were collected and immunoblotted with antibodies against phosphorylated Akt (p-Akt; S473), total Akt (T-Akt), **b** phosphorylated Erk1/2 (p-Erk1/2), total Erk1/2 (T-Erk1/2), DARPP-32 and α-tubulin (loading control). **c** A549 and H226 cells were transduced with control, DARPP-32, t-DARPP or T34A DARPP-32 overexpressing clones. Phosphorylated Akt (p-Akt; S473), total Akt (T-Akt), **d** phosphorylated Erk1/2 (p-Erk1/2), total Erk1/2 (T-Erk1/2), DARPP-32 and α-tubulin (loading control) proteins were detected by immunoblotting of cell lysates. Densitometry of the indicated immunoblots was performed using ImageJ software. All bar graphs represent mean ± SE of at least 3 independent experiments. \**P*<0.05, \*\**P*<0.01 and \*\*\**P*<0.001, one-way ANOVA.

### DARPP-32 promotes lung cancer cell migration

DARPP-32 is upregulated in various cancers including breast and gastric cancer, in which expression of DARPP-32 is associated with increased migration and invasion (Hansen et al., 2009; Zhu et al., 2013). To determine the role of DARPP-32 in NSCLC motility, we performed scratch wound healing assays using A549 and H1650 lung adenocarcinoma cells. We observed a significant decrease in cellular migration of DARPP-32 shRNA silenced A549 and H1650 cells compared to controls (Fig. 3a, b). We next investigated whether overexpression of DARPP-32 enhances cell motility in lung cancer cells. DARPP-32, as well as t-DARPP and the T34A DARPP-32 mutant, promoted increased migration in A549 and H1650 cells (Fig. 3c, d). To validate this result using a more physiologically relevant three-dimensional culture system, we performed Matrigel spot assays to assess lung tumor cell migration. A549 and H1650 lung adenocarcinoma cells stably transduced with lentivirus encoding control or DARPP-32 shRNAs were mixed with Matrigel and spotted on a cell culture plate followed by addition of medium. Similar to our previous findings, DARPP-32 knockdown substantially decreased tumor cell migration in A549 and H1650 cells (Supplementary Fig. 2a, b). Moreover, A549 and H1650 cells stably overexpressing DARPP-32, t-DARPP or mutant DARPP-32 (T34A) increased cell migration compared to control (Supplementary Fig. 3a, b). Taken together, our results suggest DARPP-32 promotes lung tumor cell migration.

**Fig. 3:**
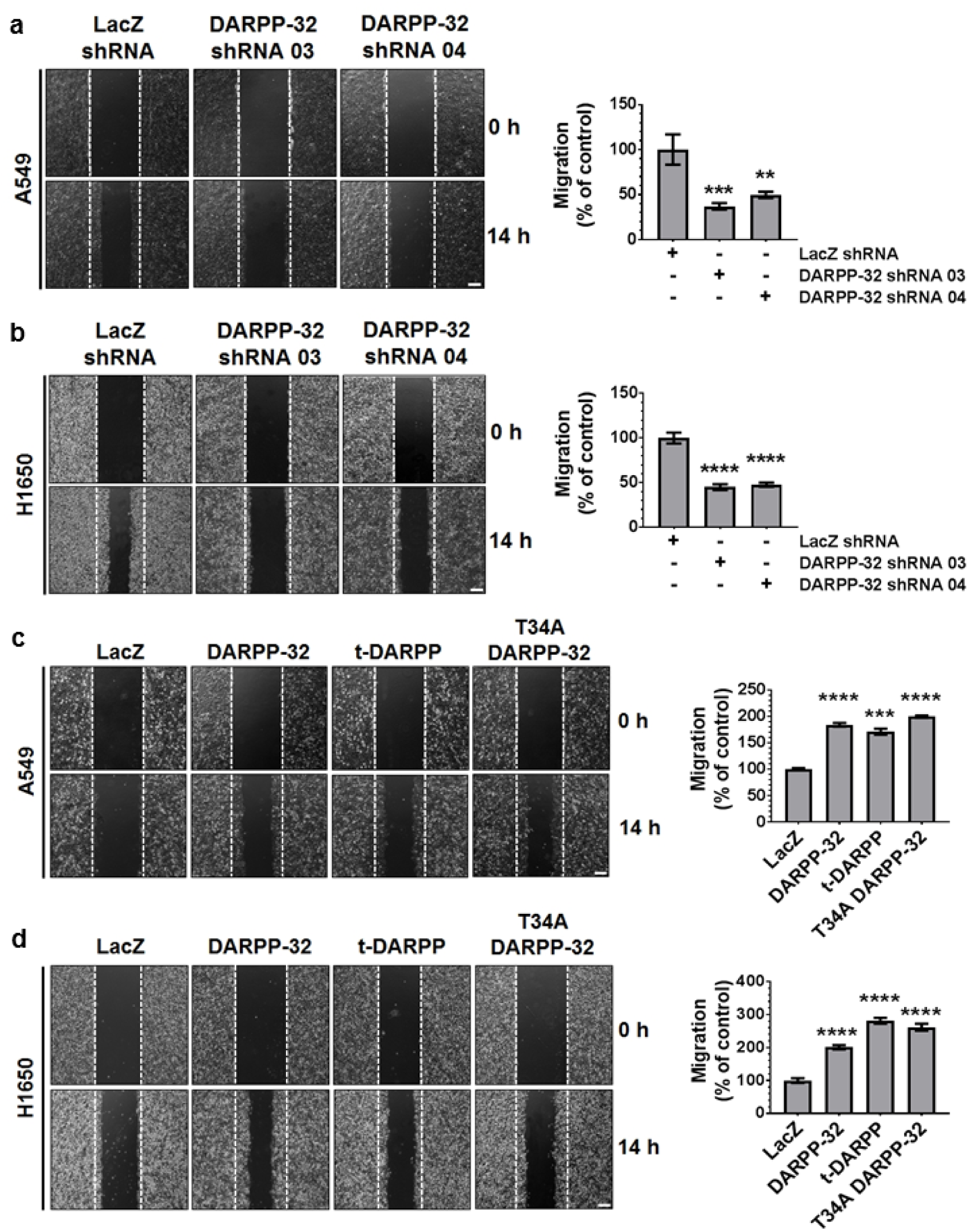
DARPP-32 positively regulate cell migration. **a** A549 and **b** H1650 cells transduced with lentivirus encoding control or DARPP-32 shRNAs were plated into 60-mm cell culture dishes, scratched and imaged at 0 and 14 h. **c** A549 and **d** H1650 cells infected with retrovirus encoding control, DARPP-32, t-DARPP or T34A DARPP-32 clones were scratched and imaged at 0 and 14 h. Dashed lines indicate the boundary of the edges of the wound at 0 h. Experiments were repeated at least three times in triplicates. Scale bar, 200 μm. Distance travelled by the migratory cells were calculated using ImageJ software. Results represent mean ± SE. \*\**P*<0.01, \*\*\**P*<0.001 and \*\*\*\**P*<0.0001, one-way ANOVA.

### DARPP-32 interacts with IKKα to activate non-canonical NF-κB2 signaling

We sought to determine the molecular signaling through which DARPP-32 promotes migration. Lung tumor cell migration has been previously shown to be regulated by non-canonical NF-κB2 signaling (Yeudall et al., 2012). Thus, we hypothesized that DARPP-32 stimulates cell migration through modulation of non-canonical NF-κB2 signaling. In stimulated cells, NF-κB inducing kinase (NIK) activates inhibitory kappa B kinase-α (IKKα), which in turn, phosphorylates cytosolic NF-κB2 p100 causing its cleavage to NF-κB2 p52, which translocates to the nucleus to transcriptionally regulate gene expression (Cildir et al., 2016). We found DARPP-32 knockdown decreased cytosolic phosphorylated NF-κB2 p100 and nuclear NF-κB2 p52 protein expression in A549 and H1650 cells (Fig. 4a, b). Given our immunoblotting result suggesting DARPP-32 promotes nuclear p52 expression, we sought to determine whether elevated DARPP-32 increases nuclear NF-κB2 p52 localization using immunofluorescence. Indeed, we observed significantly greater nuclear localization of p52 in A549 and H1650 lung cancer cells overexpressing DARPP-32, t-DARPP, or DARPP-32 T34A relative to controls (Fig. 4c, d). In accordance with these results, western blot data confirmed that overexpression of DARPP-32 activates NF-κB2 signaling by increasing the expression of cytosolic phosphorylated p100 and nuclear p52 protein expression (Supplementary Fig. 4). Interestingly, knockdown (Fig. 4a, b) or overexpression (Supplementary Fig. 4b, d) of DARPP-32 had no effect on phosphorylation of cytosolic IKKα, suggesting activation of NF-κB2 signaling is regulated by DARPP-32 in an NIK-independent manner. Thus, we sought to determine whether DARPP-32 is capable of activating NF-κB2 signaling in an NIK-independent manner through a direct interaction with IKKα. We demonstrate a physical association between DARPP-32 and IKKα through co-immunoprecipitation studies in A549 and H1650 human lung adenocarcinoma cells (Fig. 4e, f). The interaction of DARPP-32 and IKKα was significantly decreased upon DARPP-32 ablation (Fig. 4e, f). Taken together, our findings suggest that DARPP-32 activates non-canonical NF-κB2 signaling by interacting with IKKα.

**Fig. 4:**
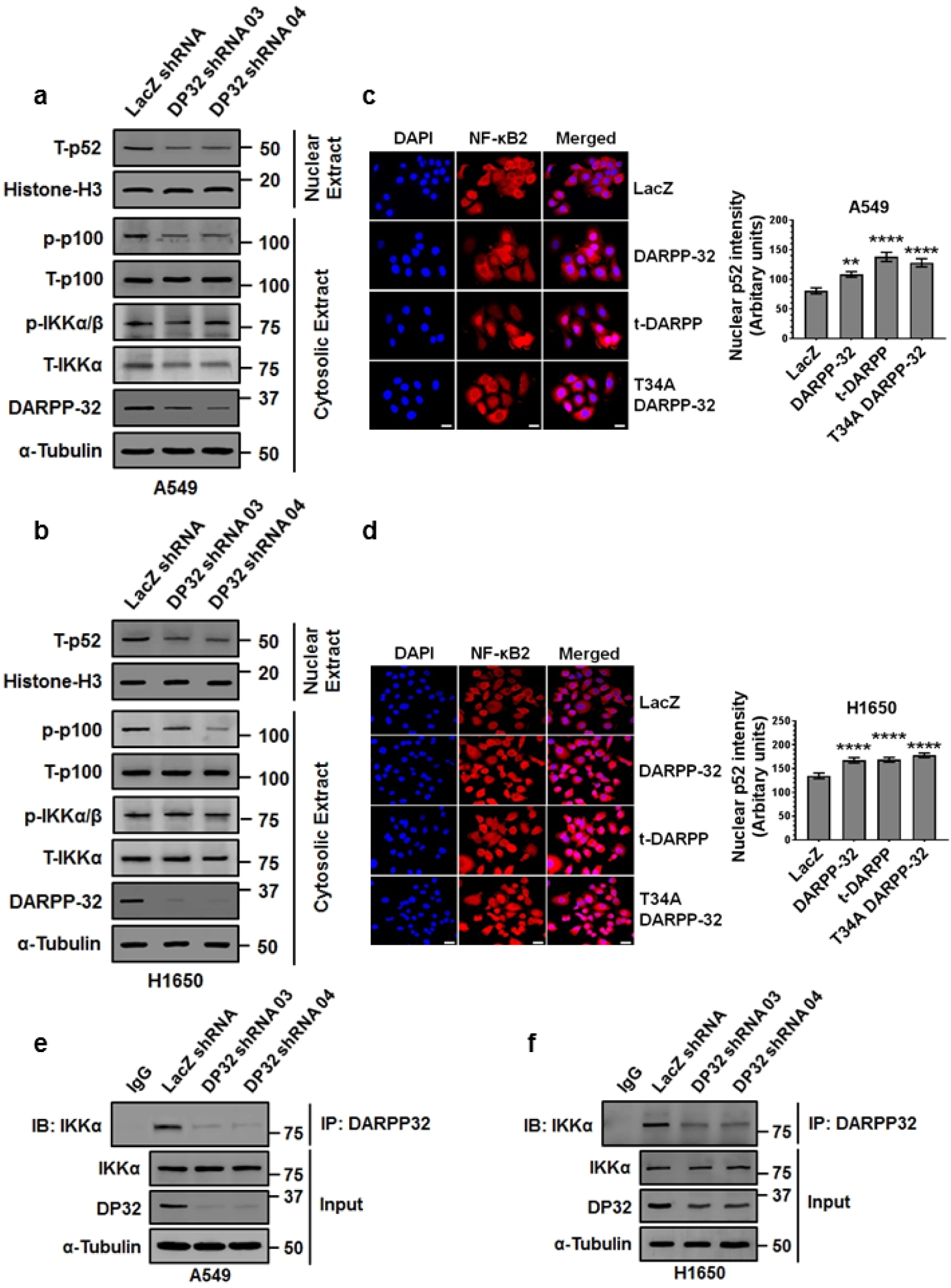
DARPP-32 knockdown inhibits non-canonical NF-κB2 signaling. **a** Nuclear fractions of A549 and **b** H1650 cells expressing control or DARPP-32 (DP32) shRNAs were immunoblotted with antibodies against total p52 (T-p52) and Histone H3 (loading control). Cytosolic fractions were also collected and subjected to western blotting using antibodies against phosphorylated p100 (p-p100), total p100 (T-p100), phosphorylated α/β (p-IKKα/β), total IKKα (T-IKKα), DARPP-32 and α-tubulin (loading control). **c** Immunofluorescence studies were performed using a monoclonal NF-κB2 antibody (that detects both p100 and p52 proteins) on A549 and **d** H1650 cell lysates expressing control or DARPP-32 shRNAs. Nuclei were labeled with DAPI. Average red fluorescence intensity of all nuclei was calculated using Image J software. Experiments were repeated at least three times. Scale bar, 20 μm. Error bars indicate SE. \*\**P*<0.01 and \*\*\*\**P*<0.0001, one-way ANOVA. **e** A549 and **f** H1650 cells transduced with lentivirus encoding control or DARPP-32 shRNAs (DP32 shRNAs) were immunoprecipitated (IP) with anti-DARPP-32 antibody and immunoblotted (IB) with antibody against IKKα. Total cell lysates (Input) were subjected to western blotting using antibodies against IKKα, DARPP-32 (DP32) and α-tubulin (loading control).

### DARPP-32 positively regulates cell migration by promoting nuclear translocation of NF-κB2 p52

We next aimed to further investigate how DARPP-32 and IKKα regulate cell migration. The IKKα-dependent non-canonical NF-κB2 pathway has a well-documented role in cell motility (Kew et al., 2012). To confirm the role of IKKα in the non-canonical NF-κB2-mediated regulation of tumor cell migration, we performed the scratch wound healing assay with control or IKKα-depleted NSCLC cells (Fig. 5a and Supplementary Fig. 5a, b). We observed a substantial decrease in migration of IKKα shRNA transduced cells compared to those expressing control shRNA (Fig. 5a). Next, we examined migration in A549 and H1650 cells upon shRNA-mediated NF-κB2 knockdown (Fig. 5b and Supplementary Fig. 5c, d). NF-κB2 depletion significantly decreased tumor cell migration in the scratch wound healing assay (Fig. 5b), suggesting both IKKα and NF-κB2 proteins are potent activators of lung tumor cell migration. Based on our cumulative findings that DARPP-32 regulates migration (Fig. 3) and activates non-canonical NF-κB2 signaling (Fig. 4), we hypothesized that DARPP-32 stimulates cell migration through IKKα and NF-κB2 signaling. To test whether DARPP-32 requires downstream IKKα and NF-κB2 signaling to promote migration, we overexpressed DARPP-32 upon shRNA-mediated knockdown of IKKα or NF-κB2 in human NSCLC cells (Supplementary Fig. 6a, b). Migration was not significantly altered in IKKα- or NF-κB2-depleted NSCLC cells upon overexpression of DARPP-32, but migration was significantly increased when DARPP-32 was overexpressed in the absence of IKKα or NF-κB2 knockdown (Fig. 5c, d). Thus, our findings suggest that DARPP-32 acts specifically through IKKα and NF-κB2 signaling to induce lung tumor migration (Supplementary Fig. 7).

**Fig. 5:**
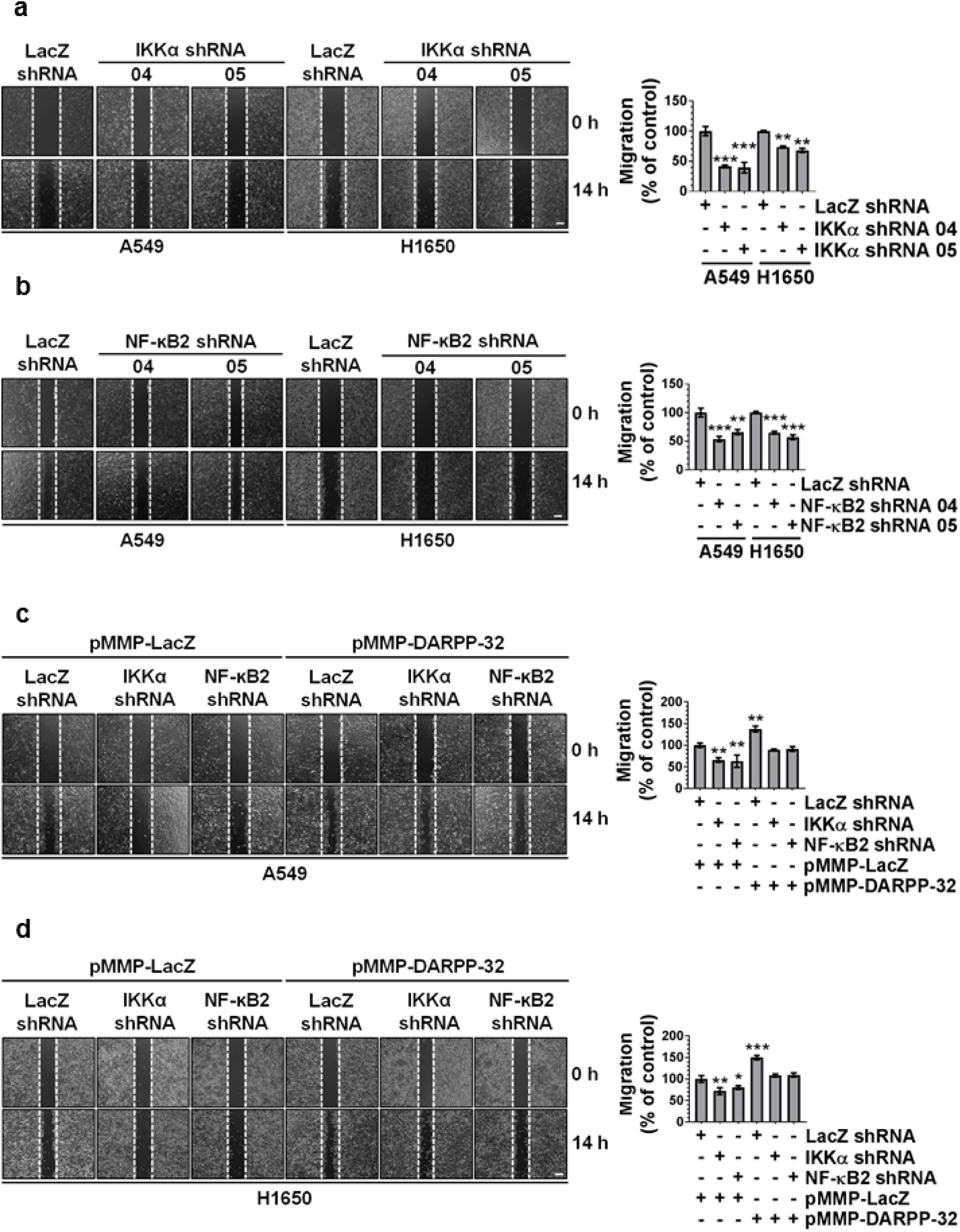
Knockdown of IKKα and NF-κB2 reduces lung cancer cell migration. **a** A549 and H1650 cells transduced with lentivirus encoding control or IKKα shRNAs (clone numbers 04 and 05) were plated into 60-mm cell culture dishes, scratched and imaged at 0 and 14 h. **b** A549 and H1650 cells transduced with lentivirus encoding control or NF-κB2 shRNAs (clone numbers 04 and 05) were scratched and imaged at 0 and 14 h. **c** A549 and **d** H1650 cells expressing control, IKKα or NF-κB2 shRNAs were transfected with control (pMMP-LacZ) or DARPP-32 (pMMP-DARPP-32) overexpressing plasmids using Polyfect reagent. The cells were then scratched and imaged at 0 and 14 h. Dashed lines indicate the boundary of the edges of the wound at 0 h. Experiments were repeated at least three times in triplicates. Scale bar, 200 μm. Distance travelled by the migratory cells were calculated using ImageJ software. Results represent mean ± SE. \**P*<0.05, \*\**P*<0.01 and \*\*\**P*<0.001, one-way ANOVA.

### DARPP-32 promotes lung tumor growth in orthotopic murine models

Based on our findings that DARPP-32 promotes lung cancer survival and migration, combined with previous studies implicating DARPP-32 as an oncogenic factor contributing to breast cancer and gastric tumor progression (Z. Chen et al., 2016; Vangamudi et al., 2010), we sought to determine whether DARPP-32 drives lung cancer growth *in vivo.* To this end, we tested whether DARPP-32 ablation reduces lung tumor growth in an orthotopic xenograft mouse model. Briefly, we injected luciferase-labeled human A549 NSCLC cells into the left thorax of anesthetized SCID mice, allowed establishment of the lung tumor and then xenogen imaged the mice regularly over the course of five to six weeks. We observed a significant decrease in lung tumor growth in mice challenged with DARPP-32 ablated A549 cells compared to mice challenged with cells transduced with control LacZ shRNA (Fig. 6a). Correspondingly, we found decreased tumor growth in mice orthotopically injected with DARPP-32 ablated H1650 (Fig. 6b) or H226 (Fig. 6c) human lung cancer cells. These findings support our hypothesis that DARPP-32 depletion inhibits human lung tumor growth. We next sought to determine whether overexpression of DARPP-32 promotes lung tumor growth *in vivo.* To address this question, we injected luciferase-labeled human A549 NSCLC cells stably overexpressing exogenous DARPP-32 or t-DARPP into the left thorax of anesthetized SCID mice. We demonstrated that DARPP-32 and t-DARPP overexpression promotes lung tumor growth in mice (Fig. 6d). Taken together, our data suggest DARPP-32 proteins drive lung tumorigenesis and inhibition of DARPP-32 reduces lung cancer growth.

**Fig. 6:**
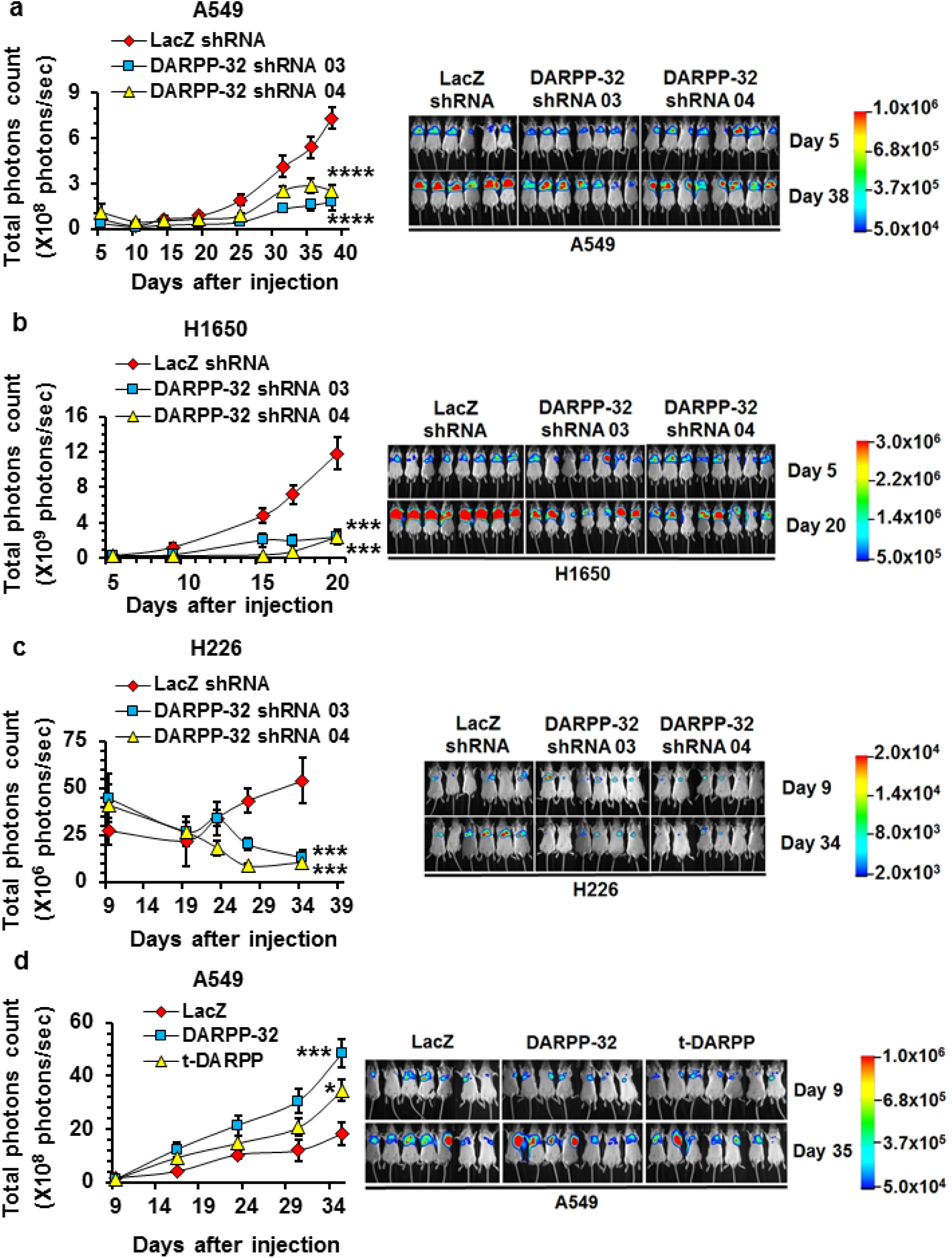
DARPP-32 knockdown decreases tumor progression in a human lung tumor xenograft model. **a** Luciferase-labeled human A549, **b** H1650 and **c** H226 cells transduced with lentivirus encoding control or DARPP-32 shRNAs were injected into the left thorax of SCID mice and imaged for luminescence on indicative days. **d** Luciferase-labeled human A549 cells overexpressing control, DARPP-32 or t-DARPP clones were orthotopically injected into the left thorax of SCID mice and imaged for luminescence on indicative days. Representative *in vivo* images of SCID mice are depicted. Total luminescence intensity (photons count) was calculated using molecular imaging software. The colored bar represents the numerical value of luminescence. Error bars indicate SE. \**P*<0.05, \*\*\**P*<0.001 and \*\*\**P*<0.0001, one-way ANOVA.

### Elevated t-DARPP expression positively correlates with tumor (T) staging score in a large cohort of lung adenocarcinoma patients

We aimed to elucidate the clinical relevance of DARPP-32 given its role in promoting tumor growth in mouse models of human NSCLC. Correspondingly, previous studies have linked upregulation of DARPP-32 and t-DARPP with breast, gastric and colorectal cancer (Z. Chen et al., 2016; Christenson et al., 2014; El-Rifai et al., 2002; Hong et al., 2012; Kopljar et al., 2015; Wang et al., 2005). To assess DARPP-32 and t-DARPP expression in NSCLC patients, we obtained tissue specimens from 62 lung adenocarcinoma patients and performed differential immunohistochemistry to detect the expression of DARPP-32 and t-DARPP. Specifically, we individually immunostained serial whole tissue sections of formalin-fixed paraffin embedded tissue blocks corresponding to each patient with two distinct DARPP-32 antibodies that: 1) detects both DARPP-32 and t-DARPP via a C-terminal epitope present in both isoforms, or 2) exclusively detects DARPP-32 through an N-terminal epitope absent in the N-terminally truncated t-DARPP isoform. Because most of the patients in our cohort had Stage III lung adenocarcinoma (Supplementary Table 1), we used the tumor (T) staging score (i.e. from the 7^th^ edition of the lung cancer TNM staging system (Mirsadraee et al., 2012)), which represents the size of the primary tumor and whether it has grown into nearby areas, as a metric of tumor progression and growth. A pulmonary pathologist (ACR) scored the percentage of positive tumor cells and their staining intensity of DARPP-32 only and both isoforms (DARPP-32 and t-DARPP) using a scale of 0-3 (i.e. 0= none, 1= weak, 2= moderate, 3= strong expression). Using the resulting pathological scoring, we calculated an immune reactive (IR) score for each specimen based on the percentage of tumor cells staining positive and the staining intensity in those cells (IR score = percentage of tumor cells x staining intensity). We found that high relative expression of t-DARPP correlates with worsening T staging score in the 62 lung adenocarcinoma specimens examined by immunohistochemistry (Fig. 7a, b). Our results suggest that a subset of patients with advanced lung adenocarcinoma exhibit elevated levels of t-DARPP protein and that upregulation of t-DARPP appears to be associated with T staging score.

**Fig. 7:**
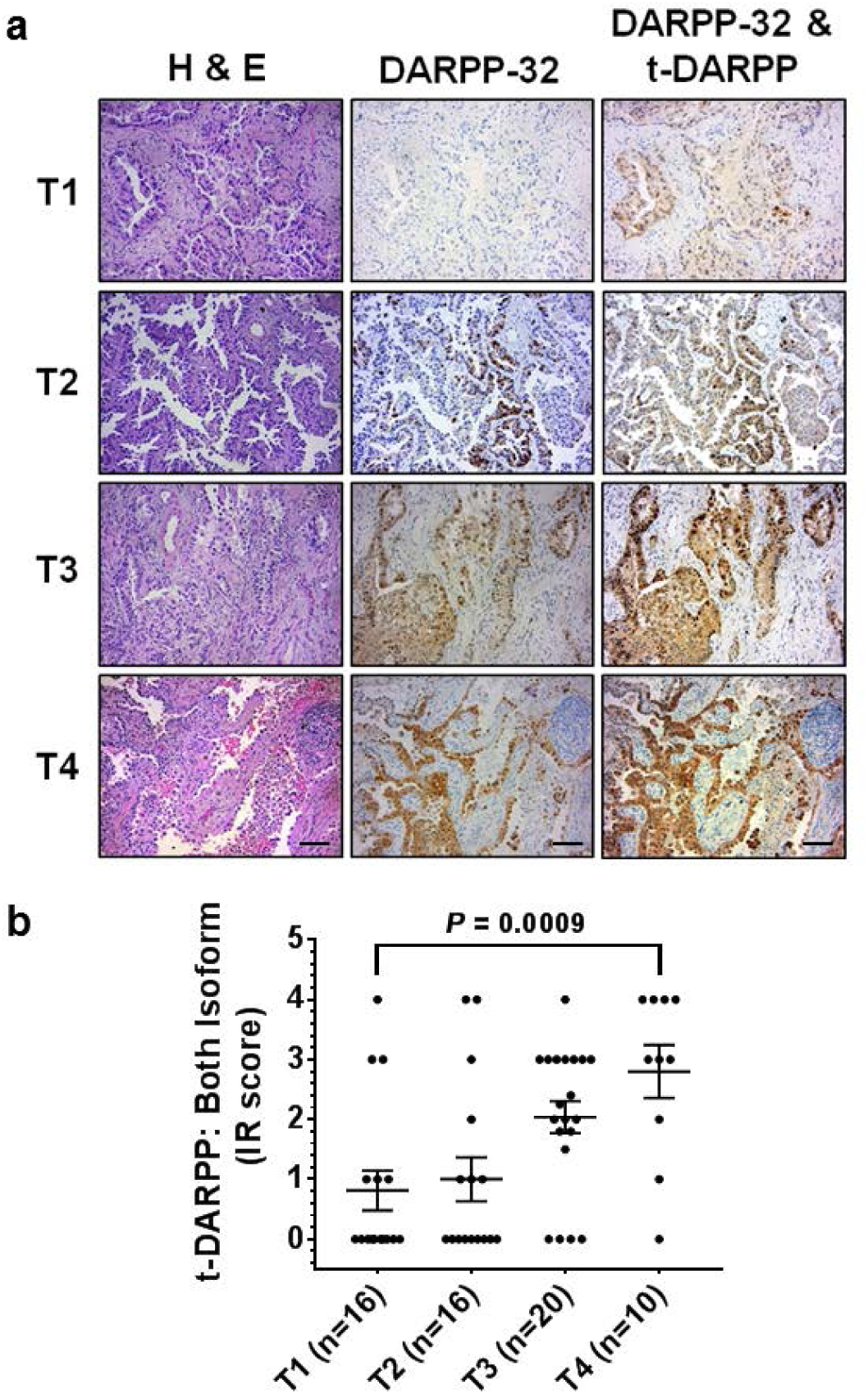
Elevated t-DARPP protein expression positively correlates with tumor (T) staging score in lung adenocarcinoma patients. **a** Immunohistochemistry was performed on human NSCLC serially sectioned specimens using an N-terminal DARPP-32 antibody to exclusively detect DARPP-32 and a C-terminal DARPP-32 antibody to detect both DARPP-32 and t-DARPP. Scale bar, 50 μm. **b** Differential immunohistochemistry was used to quantify t-DARPP protein expression in 62 human lung cancer tissue samples. Each tissue was scored based on the percentage of tumor cells stained positive multiplied by the staining intensity (i.e. 0= none, 1= weak, 2= moderate, 3= strong expression) to yield an immune reactive (IR) score. The IR score for t-DARPP protein expression was calculated by subtracting the IR score of DARPP-32 (detected with N-terminal antibody) from the IR score of both DARPP-32 isoforms (detected with C-terminal antibody). Each point on the graph represents an individual tissue. Error bars indicate SE.

### Upregulation of t-DARPP, NF-κB2 and IKKα in lung adenocarcinoma is associated with decreased patient survival

We utilized a bioinformatics approach to validate our finding that high relative t-DARPP expression correlates with tumor growth in lung adenocarcinoma patients. We assessed relative DARPP-32 and t-DARPP transcript expression in specimens corresponding to 513 human lung adenocarcinoma patients cataloged in The Cancer Genome Atlas (TCGA). Interestingly, we found that expression of t-DARPP increases with advancing tumor (T) stages in lung adenocarcinoma (Fig. 8a). As assessed by KaplanMeier survival curve, we observed that patients with high t-DARPP expression showed substantially decreased survival relative to lung adenocarcinoma patients with low t-DARPP expression (Fig. 8b). Our findings indicate that t-DARPP expression is an important determinant of survival in lung adenocarcinoma patients.

**Fig. 8:**
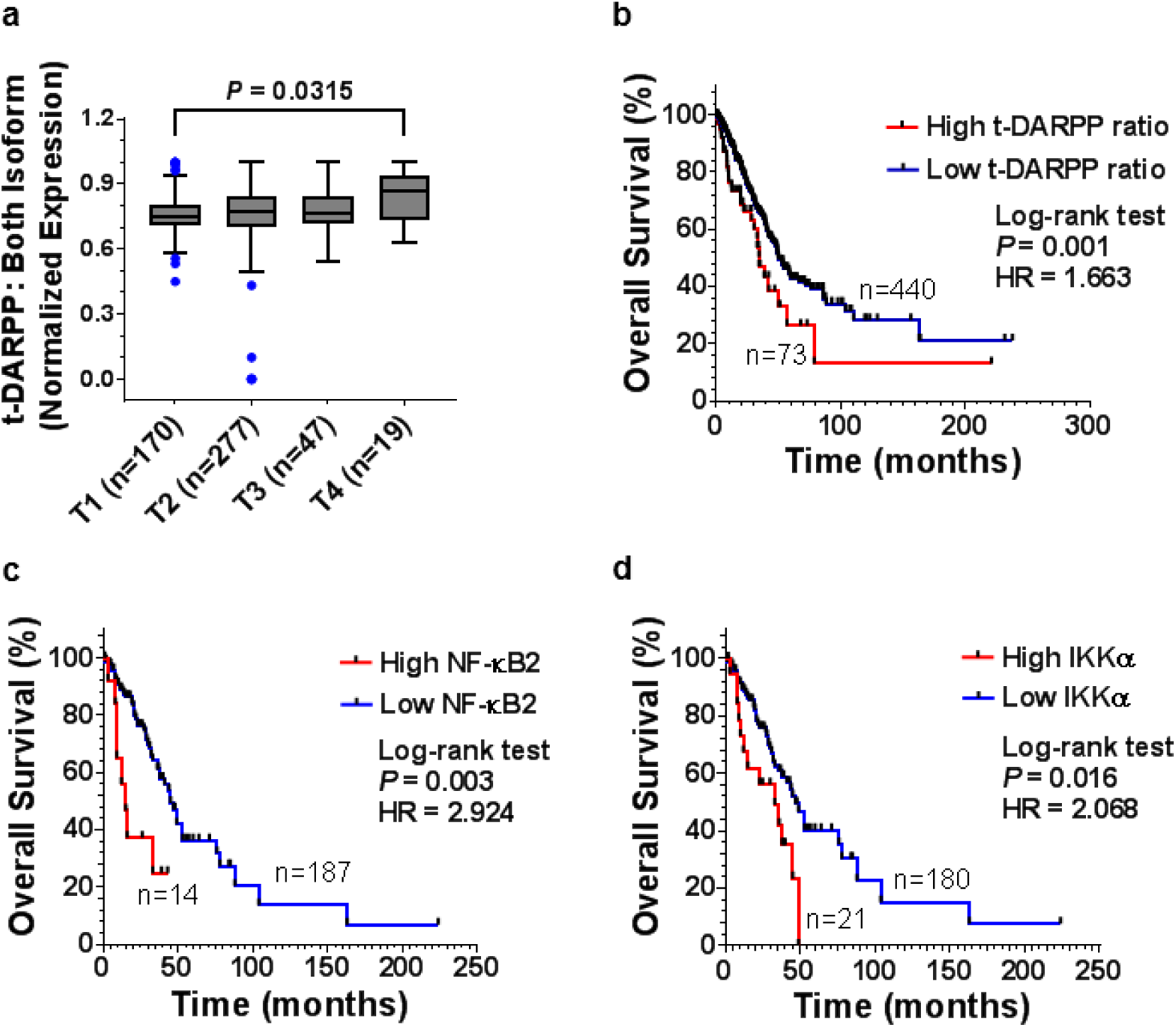
Elevated t-DARPP, NF-κB2 and IKKα transcripts correlate with decreased lung adenocarcinoma patient survival. **a** Quantification of t-DARPP mRNA expression in 513 human lung cancer tissue samples. Blue dots indicate outliers. T1-T4 represents tumor (T) staging scores from TNM system. **b** Kaplan Meier plot showing overall survival within the total cohort of 503 NSCLC patients based on the expression of t-DARPP and DARPP-32 mRNAs. **c** Kaplan Meier curve depicting overall survival within the total cohort of 201 NSCLC patients based on the expression of NF-κB2 or **d** IKKα mRNAs. The normalized read count for DARPP-32, t-DARPP, NF-κB2 and IKKα mRNAs were obtained from The Cancer Genome Atlas dataset (TCGA). The difference between the two groups was calculated using the Log-rank (Mankel-Cox) test. HR: hazard ratio.

Given our findings that DARPP-32 isoforms regulate non-canonical NF-κB2-mediated cell migration, we asked whether expression of NF-κB2 or IKKα is associated with overall survival of lung adenocarcinoma patients. RNA-Seq expression data from 201 human lung adenocarcinoma tissue samples was used to generate Kaplan-Meier survival curves. Our results reveal significantly decreased survival in the patients with high expression of NF-κB2 and IKKα transcripts compared to low expressers of those mRNAs (Fig. 8c, d). Thus, upregulation of NF-κB2 and IKKα expression is associated with decreased overall patient survival and may predict poor clinical outcome in lung adenocarcinoma patients.

## Discussion

For the first time, we demonstrate DARPP-32 and its splice variant t-DARPP stimulate lung cancer cell survival and migration to promote oncogenesis, and we show elevated t-DARPP isoform levels in NSCLC patients are associated with increased tumor staging and worsened patient survival. The role of DARPP-32 and t-DARPP in cancer has emerged beyond their classical function as modulators of dopamine-mediated neurotransmission, highlighting their importance in the regulation of physiological and pathological effects. For example, alternations in expression of DARPP-32 and t-DARPP have been implicated in schizophrenia, bipolar disorder and Alzheimer’s disease (Cho et al., 2015; Kunii et al., 2014) as well as numerous types of tumors, including breast, gastric, prostate, esophageal and colon cancers (Beckler et al., 2003; El-Rifai et al., 2002; Hamel et al., 2010; Vangamudi et al., 2010). Since investigation of the frequent amplification at the 17q12 locus in gastric cancers implicated DARPP-32 and t-DARPP in oncogenesis (Belkhiri et al., 2005; El-Rifai et al., 2002), numerous studies have demonstrated the role of these proteins in cancer cell survival, drug resistance, migration, invasion and angiogenesis (Belkhiri et al., 2016).

Our results suggest DARPP-32 and t-DARPP promote NSCLC cell survival through activation of Akt and Erk1/2 signaling by protecting cells from apoptotic cell death (Figs. 1–2). Correspondingly, overexpression of DARPP-32 and t-DARPP in human gastrointestinal adenocarcinoma cells was shown to cause a four-fold reduction in apoptosis (Belkhiri et al., 2005). The T75 phosphorylation residue shared by DARPP-32 and t-DARPP was attributed to promoting cell survival (Belkhiri et al., 2005), and a follow-up report by the same group suggested increased activation of Akt and Bcl2 is mechanistically responsible for t-DARPP-mediated cancer cell survival (Belkhiri, Dar, Zaika, et al., 2008). Evasion of apoptosis is a major underlying mechanism in the ability of cancer cells to acquire resistance to molecular targeted therapies (Rotow et al., 2017). El-Rifai and colleagues demonstrated that DARPP-32 promotes cell survival and gefitinib resistance in gastric cancer cells by stimulating EGFR phosphorylation and activating PI3K/Akt signaling (Zhu et al., 2011). DARPP-32 stimulated resistance to pro-apoptotic proteins through induction of pro-survival molecule Bcl-xL through Src/STAT3 signaling cascades (Belkhiri et al., 2012). Numerous reports have implicated t-DARPP in breast cancer patients acquiring resistance to trastuzumab (Herceptin), a monoclonal antibody targeting the ERBB2 (Her2/neu) receptor. Collectively, these studies demonstrated that t-DARPP drives breast cancer cell resistance to trastuzumab through inhibition of apoptotic caspase-3 and activation of pro-survival Akt signaling through its T75 residue, common among both DARPP-32 isoforms (Belkhiri, Dar, Peng, et al., 2008; Gu et al., 2009; Hamel et al., 2010). Another report showed t-DARPP promotes trastuzumab resistance in esophageal adenocarcinoma cells through similar mechanisms (Hong et al., 2012). In both breast and esophageal cancer, t-DARPP physically interacted with ERBB2 in a protein complex to mediate trastuzumab resistance (Belkhiri, Dar, Peng, et al., 2008; Hong et al., 2012). Like previous studies in gastric, breast and esophageal cancers, our studies suggest DARPP-32 and t-DARPP promote cell survival through upregulation of Akt signaling. Despite this observation and the common underlying mechanistic evidence, future studies beyond the scope of this manuscript are necessary to determine whether t-DARPP promotes resistance to specific molecular targeted NSCLC therapies.

Like pro-survival mechanisms, increased cell migration also contributes to cancer cell growth and resistance to molecular targeted therapies. Given the well-established association between DARPP-32 isoforms and acquired drug resistance in cancer, it is unsurprising that several reports and detailed reviews have described the role of DARPP-32 in breast and gastric cancer cell migration and invasion (Belkhiri et al., 2016; Hansen et al., 2006; Zhu et al., 2013). Correspondingly, we provide evidence that DARPP-32 and t-DARPP promote NSCLC cell migration based on *in vitro* scratch (Fig. 3) and spot assays (Supplementary Figs. 2 and 3). In line with our observations, other studies have demonstrated that overexpression of DARPP-32 promotes tumor cell invasion in gastric and colorectal cancers (Kopljar et al., 2015; Zhu et al., 2013). Conversely, DARPP-32 has been shown to inhibit breast cancer cell migration through a dopamine D1 receptor-dependent mechanism (Hansen et al., 2006). A subsequent *in vitro* study has revealed that PP-1 inhibition regulated by phosphorylation of DARPP-32 at residue T34 is critical for modulating cell migration in breast cancer (Hansen et al., 2009). Taken together, the regulation of cancer cell migration by DARPP-32 is likely cell and tumor type dependent.

We identify a novel physical interaction between DARPP-32 and IKKα that suggests DARPP-32 regulates non-canonical NF-κB2 signaling to control NSCLC migration (Fig. 4). Knockdown of IKKα, as well as independently silencing NF-κB2, decreased migration of human lung adenocarcinoma cells (Fig. 5a, b). Based on our findings, we propose that DARPP-32 activates IKKα through an unknown NIK-independent mechanism that leads to IKKα-mediated phosphorylation of NF-κB2 p100, ubiquitination and partial degradation of p100 to p52, and translocation of NF-κB2 p52 to the nucleus where it acts as a transcription factor to modulate expression of genes involved in cell migration (Supplementary Fig. 7). A recent report has demonstrated that *Helicobacter pylori* infection induces canonical NF-κB1-mediated transcriptional upregulation of DARPP-32 mRNA and protein, which counteracts *Helicobacter pylori*-mediated cell death through activation of Akt (Zhu, Soutto, et al., 2016). Therefore, we investigated whether non-canonical NF-κB2 signaling altered DARPP-32 protein expression, but observed no effect (Supplementary Fig. 5). While canonical NF-κB1 pathway activation has been linked to the growth and survival of many solid and hematological malignancies, the role of non-canonical NF-κB2 signaling in cancer is still emerging (Cildir et al., 2016; Hayden et al., 2008). However, studies suggest the NF-κB2 pathway is activated in cancer through viral oncogenes, mutations in pathway components, and upregulation of upstream components of the pathway (Cildir et al., 2016), the latter of which is supported by our results, suggesting DARPP-32 promotes activation of non-canonical NF-κB2 signaling in lung cancer through an interaction with IKKα.

We demonstrate stable overexpression of DARPP-32 and t-DARPP in human NSCLC cells orthotopically implanted into the thoracic cavity of SCID mice promotes tumor growth (Fig. 6d). Correspondingly, mice that received an orthotopic xenograft of shRNA-mediated DARPP-32 silenced NSCLC cells exhibited decreased tumor growth relative to controls (Fig. 6a, b, c). El-Rifai and colleagues have shown overexpression of t-DARPP in human OE19 esophageal adenocarcinoma subcutaneously xenografted into athymic nude mice stimulates tumor growth (Hong et al., 2012). Using a similar xenograft mouse model, they have subsequently demonstrated shRNA-mediated knockdown of DARPP-32 reduces gastric tumorigenesis (Zhu et al., 2011; Zhu, Chen, et al., 2016) and overexpression of DARPP-32 in AGS human gastric adenocarcinoma cells promotes *in vivo* tumor growth (Z. Chen et al., 2016). To the best of our knowledge, our study is the first to assess DARPP-32 knockdown as well as DARPP-32 and t-DARPP overexpression in an orthotopic cancer xenograft mouse model. Importantly, our *in vivo* results showing DARPP-32 and t-DARPP promote NSCLC oncogenesis coincide with similar findings in esophageal and gastric cancer subcutaneous xenograft models.

Based on differential immunostaining of over 60 human NSCLC specimens, we describe that high relative expression of t-DARPP correlates with tumor staging in lung adenocarcinoma patients (Fig. 7). Similar differential immunohistochemistry approaches in serial tissue sections have been previously used to distinguish between detection of DARPP-32 only (N-terminal antibody) versus both isoforms (C-terminal antibody). Two independent studies have demonstrated a subset of primary human breast cancer specimens exhibit elevated t-DARPP protein levels relative to DARPP-32 (Hamel et al., 2010; Vangamudi et al., 2010). Using a genetic spontaneous murine model of breast cancer, Christenson and Kane have found DARPP-32 was expressed in normal mammary tissue and in some breast tumors, whereas t-DARPP was detected exclusively in tumors, typically at higher or equal levels as DARPP-32 (Christenson et al., 2014). This transition from DARPP-32 to t-DARPP observed during breast tumorigenesis corresponds to our pathological and bioinformatics findings linking upregulation of t-DARPP expression with increased NSCLC growth and worsened patient survival. The DARPP-32 to t-DARPP isoform shift in cancer may be directed by the SRp20 splicing factor, which has been shown to physically associate with DARPP-32 (Zhu, Chen, et al., 2016). The upregulation of t-DARPP in NSCLC progression suggests its expression stimulates oncogenesis. Thus, t-DARPP may represent a promising molecular target in NSCLC as well as possess prognostic value.

## Methods

### Cell Culture

Human NSCLC cell lines A549, H1650 and H226 as well as the transformed human embryonic kidney epithelial cell line, HEK-293T, were purchased from American Type Culture Collection (Manassas, VA) and maintained according to the manufacturer’s instructions. HEK-293T cells were cultured in Dulbecco’s modified Eagle’s medium (DMEM; Corning; Manassas, VA) and lung cancer cell lines were cultured in RPMI-1640 medium (Corning). Media was supplemented with 10% fetal bovine serum (FBS; Millipore; Burlington, MA) and 1% Penicillin/Streptomycin antibiotics (Corning). All cell lines were certified by the indicated cell bank and routinely authenticated by morphologic inspection.

### Transient transfections

5×10^5^ A549 or H1650 cells were seeded in 60-mm cell culture plates and incubated for 24 h in RPMI-1640 medium. Cells were then washed with PBS, suspended in OPTI-MEM reduced serum medium (Gibco; Grand Island, NY), and transfected with 2 μg of pMMP-LacZ or pMMP-DARPP-32 plasmids using Polyfect transfection reagent (Qiagen; Hilden, Germany) according to instruction from the manufacturer. After 4 h, antibiotic-containing complete RPMI-1640 medium was added and cells were grown until they had established a confluent monolayer.

### Generation of stable cell lines

Expression constructs of human DARPP-32, t-DARPP and DARPP-32 T34A cDNA in pcDNA3.1 were a generous gift from Dr. Wael El-Rifai at Vanderbilt University Medical Center (Z. Chen et al., 2016). The Flag-tagged coding sequence of DARPP-32, t-DARPP and T34A DARPP-32 were subcloned into the retroviral (pMMP) vector. The pMMP plasmid and its corresponding pMMP-LacZ control construct were kindly provided by Dr. Debabrata Mukhopadhyay at Mayo Clinic in Jacksonville, Florida (Zeng et al., 2001). Production of retrovirus and transduction of A549, H1650 and H226 lung cancer cell lines were performed as previously described (Zeng et al., 2001).

Four to five different lentiviral shRNA pLKO.1 constructs (Sigma-Aldrich; St. Louis, MO) were used to silence protein expression of each target, including DARPP-32, NF-κB2 and IKKα. pLKO.1-LacZ shRNA (Sigma-Aldrich) was used as a corresponding control. Generation of lentivirus and transduction of A549, H1650 and H226 lung cancer cell lines were accomplished as previously described (Alam et al., 2016).

### Cell survival assay

A549, H226 and H1650 human NSCLC cell lines were each plated in a 96-well microplate at a concentration of 3000 cells/well. Cell viability was assessed after 72 h of incubation using CellTiter 96^®^ AQueous One System (Promega; Madison, WI). Absorbance was recorded at 490 nm using an Epoch microplate spectrophotometer (Biotek; Winooski, VT). The average of three independent experiments has been reported.

### Cell proliferation analysis by BrdU labeling

Human NSCLC cells were seeded at a density of 1×10^5^ cells per 60-mm plate. The following day, bromodeoxyuridine (BrdU; 30 μM; Sigma-Aldrich) diluted in fresh medium was administered to the cells for 30 minutes. The cells were harvested, fixed, and processed for incubation with primary mouse anti-BrdU monoclonal antibody (Roche; Madison, WI) and subsequently secondary APC-conjugated goat anti-mouse antibody (Biotium; Fremont, CA). BrdU-positive cells were counted. Finally, the cells were stained with propidium iodide (Sigma-Aldrich) for flow cytometry analysis. The average of three separate experiments has been documented.

### Apoptosis analysis

1×10^5^ LacZ control- or DARPP-32 shRNA-transduced A549, H1650 and H226 cells were plated in 60mm dishes. Cells were then harvested, washed with PBS, and stained with Annexin V-APC (BD Biosciences; San Jose, CA) following 24h incubation. Additional exposure to propidium iodide (BD Biosciences) made it possible to differentiate early apoptotic cells (Annexin-positive and propidium-iodide-negative) from late apoptotic cells (Annexin-positive and propidium-iodide-positive). Apoptotic cell death was assessed by counting the number of cells stained positive for Annexin V-APC. The average of three independent experiments has been reported.

### Antibodies

Antibodies against DARPP-32 (Cat no.:sc-11365) and α-Tubulin (Cat no.:sc-5286) were purchased from Santa Cruz Biotechnology (Dallas, TX). Antibodies against phosphorylated Akt (S473; Cat no.:4060), total Akt (Cat no.:4691), phosphorylated p44/42 MAPK (T202/Y204; Cat no.:4370), total p44/42 MAPK (Cat no.:4695), phosphorylated NF-κB2 p100 (S866/870; Cat no.:4810), total NF-κB2 p100/52 (Cat no.:4882), phosphorylated IKKα/β (S176/180; Cat no.:2697), total IKKα (Cat no.: 11930), histone H3 (Cat no.:4499) and FLAG tag (Cat no.:14793) were obtained from Cell Signaling Technology (Danvers, MA). An antibody that extensively detects DARPP-32 (Cat no.:ab40801) via an N-terminal epitope absent in t-DARPP was purchased from Abcam (Cambridge, MA) and used for immunohistochemistry. Horseradish peroxidase (HRP)-conjugated anti-rabbit (Cat no.:7074) and anti-mouse (Cat no.:7076) secondary antibodies were also purchased from Cell Signaling Technology.

### Nuclear extract preparation

5×10^6^ human NSCLC cells were suspended in hypotonic buffer [20 mM Tris-Cl pH 7.4, 10 mM NaCl, 3 mM MgCl_2_, protease inhibitor cocktail, and 1 mM phenylmethylsulfonyl fluoride (PMSF; Cell Signaling Technology)] and incubated on ice for 15 minutes. Nonionic detergent NP-40 (10%; Sigma Aldrich) was then added to the cell suspension, which was mixed vigorously. Next, the cell homogenate was centrifuged at 5,000 rpm for 10 minutes at 4°C. The supernatant was collected as the cytoplasmic fraction, and the pellet was suspended in cell extraction buffer (Thermo Fisher Scientific; Waltham, MA) supplemented with protease inhibitor cocktail (Roche) and 1 mM PMSF. The suspension was incubated on ice for 30 minutes with intermittent vortexing. Finally, the sample was centrifuged at 14,000 g for 30 minutes at 4°C, and the supernatant was collected as nuclear extract.

### Immunoblotting

A549, H1650 and H226 cells were sonicated and lysed in RIPA buffer (Millipore) containing protease inhibitor cocktail (Roche). Proteins were separated via 4-20% gradient SDS-PAGE (Bio-Rad; Hercules, CA) and transferred to polyvinyl difluoride membranes (PVDF; Millipore). Membranes were blocked with 5% bovine serum albumin (BSA; Sigma-Aldrich) and incubated with primary and secondary antibodies. Antibody-reactive protein bands were detected by enzyme-linked chemiluminescence (Thermo Fisher Scientific).

### Immunoprecipitation

Human lung cancer cells were homogenized and lysed in RIPA buffer (Millipore) supplemented with protease inhibitor cocktail (Roche). Protein concentration was measured using the Quick Start Bradford protein assay (Bio-Rad) and 500 μg of protein lysate was loaded into the supplied spin column (Catch and Release Immunoprecipitation Kit; Millipore). Immunoprecipitation was achieved by following manufacturer’s protocol (Cat no.:17-500; Millipore).

### Immunofluorescence

A549, H1650 and H226 cells were fixed in 4% paraformaldehyde (Boston Bioproducts; Ashland, MA), permeabilized in cold methanol (Fisher Scientific; Hampton, NH), and immunofluorescence staining was performed using an antibody against NF-κB2 p100/52 (Cell Signaling Technology; Cat no.:3017). The cells were then washed with PBS and incubated with Alexa Fluor 568-conjugated anti-rabbit secondary antibody (Molecular Probes; Eugene, OR; Cat no.:A11036). All images were captured by confocal microscopy (Nikon; Minato, Tokyo, Japan; 60x objective, 1.27 NA) and processed using ImageJ software (Version 1.6.0_24; https://imagej.nih.gov/ij). DAPI-stained nuclei were segmented manually by drawing a circular ROI. Nuclear colocalization of NF-κB2 p52 (red fluorescent signal) with DAPI was measured. The mean nuclear fluorescence was calculated and plotted in GraphPad Prism software (Version 7).

### Immunohistochemistry

Human lung adenocarcinoma tissue specimens were obtained from 62 NSCLC patients at Mayo Clinic in Rochester, MN in accordance with IRB approved protocols. We performed differential immunohistochemistry using an N-terminal antibody that extensively recognizes DARPP-32 (Abcam; Cat No.:ab40801). We used another C-terminal antibody that recognizes both DARPP-32 and t-DARPP (Santa Cruz Biotechnology; Cat No.:sc-11365;). Formalin-fixed, paraffin-embedded whole tissues were serially sectioned and immunostained for DARPP-32 using a Bond Autostainer (Leica; Wetzlar, Germany) as previously described (Hoeppner et al., 2015). H&E staining was also performed. In each lung tumor specimen, the intensity and prevalence of DARPP-32 staining in various cell types was scored by a pulmonary pathologist (ACR).

### Scratch wound assay

A549 and H1650 cells were seeded in 60-mm culture dishes at an appropriate density to achieve a confluent monolayer. After 16h, a linear scratch wound was generated using a sterile 20 μl pipette tip. Cells were imaged at time 0 and 14h post-scratch induction. All the images were captured using a 4X Plan S-Apo 0.16 NA objective on an EVOS FL cell imaging system (Thermo Fisher Scientific). The images were analyzed using ImageJ software and cell migration was quantified as previously described (Liang et al., 2007).

### Spot Assay

A549 and H1650 human lung cancer cells were trypsinized and suspended in RPMI-1640 medium (Corning) at a concentration of 5×10^4^ cells per μl. Cells (2.5×10^5^ in 5 μl) were then mixed with Matrigel^®^ Basement Membrane Matrix (Corning) in 1:1 ratio and pipetted as a spot in a 60-mm culture dish. Matrigel containing cell suspension (i.e. the spot) was allowed to solidify by incubating at 37°C for 5 minutes. Thereafter, medium was added and images were captured using a 4X Plan S-Apo 0.16 NA objective on an EVOS FL cell imaging system (Thermo Fisher Scientific). After a 96h incubation, the spots were imaged again and cell migration was calculated as previously described (Kaur et al., 2012).

### *In vivo* orthotopic lung cancer model

Six to eight-week-old pathogen-free SCID/NCr mice were purchased from the Charles River Laboratories. Mice were allowed one week to acclimate to their surroundings, maintained under specific pathogen-free conditions in a temperature-controlled room with alternating 12h light/dark cycles and fed a standard diet. Mice were orthotopically injected with 1×10^6^ luciferase-labeled human A549, H226 and H1650 lung cancer cells suspended in 80 μl PBS and Matrigel. After establishment of the lung tumor, mice were imaged using an In-Vivo Xtreme xenogen imaging system (Bruker; Billerica, MA) to measure luciferase intensity. To determine tumor growth, luciferase intensity was calculated using Bruker molecular imaging software and plotted over time in GraphPad Prism 7 software. All animal studies were performed in accordance with protocols approved by the University of Minnesota Institutional Animal Care and Use Committee.

### RNA-Seq analysis

The RNA-Seq isoform expression data of human lung adenocarcinoma tissue specimens in The Cancer Genome Atlas (TCGA) were used for correlation analysis with the clinical variables. The clinical information of 513 patients were downloaded from the Xena Public Data Hubs (https://xena.ucsc.edu). The processed RNA-Seq data (version 2 Level 3) for the normalized isoform expression of the 513 tumor samples were downloaded from the Genomic Data Commons Legacy Archive (https://gdc-portal.nci.nih.gov/legacy-archive). DARPP-32 contains five isoforms in the downloaded data. Among the five isoforms, uc002hrz.2, uc002hsa.2, uc010cvx.2 are longer isoforms (i.e. representing full length DARPP-32) starting at the original start codon, while uc002hsb.2 and uc002hsc.2 are shorter isoforms (i.e. representing alternate isoform t-DARPP) sharing another downstream start codon. The log2 (x+1) transformed FPKM (fragments per kilobase of transcript per million mapped reads) value normalized by RSEM (reads per kilobase of transcript per million mapped reads) for isoform and gene expressions were used in further application. Based on the t-DARPP expression, a Kaplan-Meier survival graph was created by using GraphPad Prism 7 software.

The NF-κB2 and IKKα expression data and the clinical variables of 203 human lung adenocarcinoma tissue specimens were obtained from cBioPortal for Cancer Genomics (http://cbioportal.org) (Cerami et al., 2012; Gao et al., 2013). Patients were categorized into 2 separate groups based on the mRNA expressions (normalized read count) and Kaplan-Meier survival curve was generated by using GraphPad Prism 7 software.

### Statistics

Data are expressed as mean ± SEM and representative of at least three independent experiments. Statistical significance was determined using one-way analysis of variance (ANOVA) and a value of *P* < 0.05 was considered significant.

## Acknowledgements

This work was supported by The Hormel Foundation, Austin “Paint the Town Pink”, National Institutes of Health, National Cancer Institute R00-CA187035 funding to L.H.H and K12-CA090628 to A.S.M. The Hormel Institute SURE program supported E.N.D. and A.M.C. We thank Dr. Debabrata Mukhopadhyay at Mayo Clinic in Jacksonville, FL for his valuable support and contributions to the development of this project, including sharing reagents and advice. We appreciate the contributions of Todd Schuster, the shared facilities manager at The Hormel Institute, to flow cytometry-based apoptosis and BrdU assays. We thank Kim Klukas and The Hormel Institute animal facilities staff for providing excellent animal care and husbandry. We are grateful to Shirley Bradley and Sandip Suresh for their research contributions to this project as a rotating graduate student and a summer undergraduate intern, respectively, at Mayo Clinic. We thank Dr. Wael El-Rifai at Vanderbilt University in Nashville, TN for sharing DARPP-32 reagents and Dr. Yasuhiro Ikeda at Mayo Clinic in Rochester, MN for providing constructs to stably luciferase label lung cancer cells. We thank the Mayo Clinic Pathology Research Core for assistance with immunohistochemistry staining and The Hormel Institute for administrative and institutional support.

## Author contributions

S.K.A. transduced the human lung cancer cells, performed *in vitro* experiments, and conducted the mouse studies. S.K.A. and M.A. performed mouse necropsy. P.L. and D.K.H. performed initial studies validating original hypotheses and optimized transfection/transduction efficacies. S.K.A., M.A., P.L., S.R.H., A.M.C., E.N.D., and D.K.H. contributed to numerous technical aspects of the project and implementation of experiments. S.K.A., M.A. and A.M.C. performed imaging for immunofluorescence and/or immunohistochemistry studies. W.Z. and R.K. performed t-DARPP and DARPP-32 bioinformatics analysis. S.K.A. used computational biology to analyze correlation of NF-κB2 and IKKα transcript expression with survival of lung adenocarcinoma patients. A.C.R. pathologically reviewed and scored immunostained human NSCLC patient specimens. A.S.M. directed the acquisition of NSCLC patient tissue and provided consultation as a thoracic oncologist. S.K.A. and L.H.H. reviewed relevant literature to intellectually advance the project, designed studies, performed experimental troubleshooting, analyzed data, generated figures, and wrote the manuscript. L.H.H. developed the original aims of this study, directed all research efforts, supervised the project, and obtained funding to support the work. All authors approved the final version of this manuscript.

---

**Supplementary Fig. 1:**
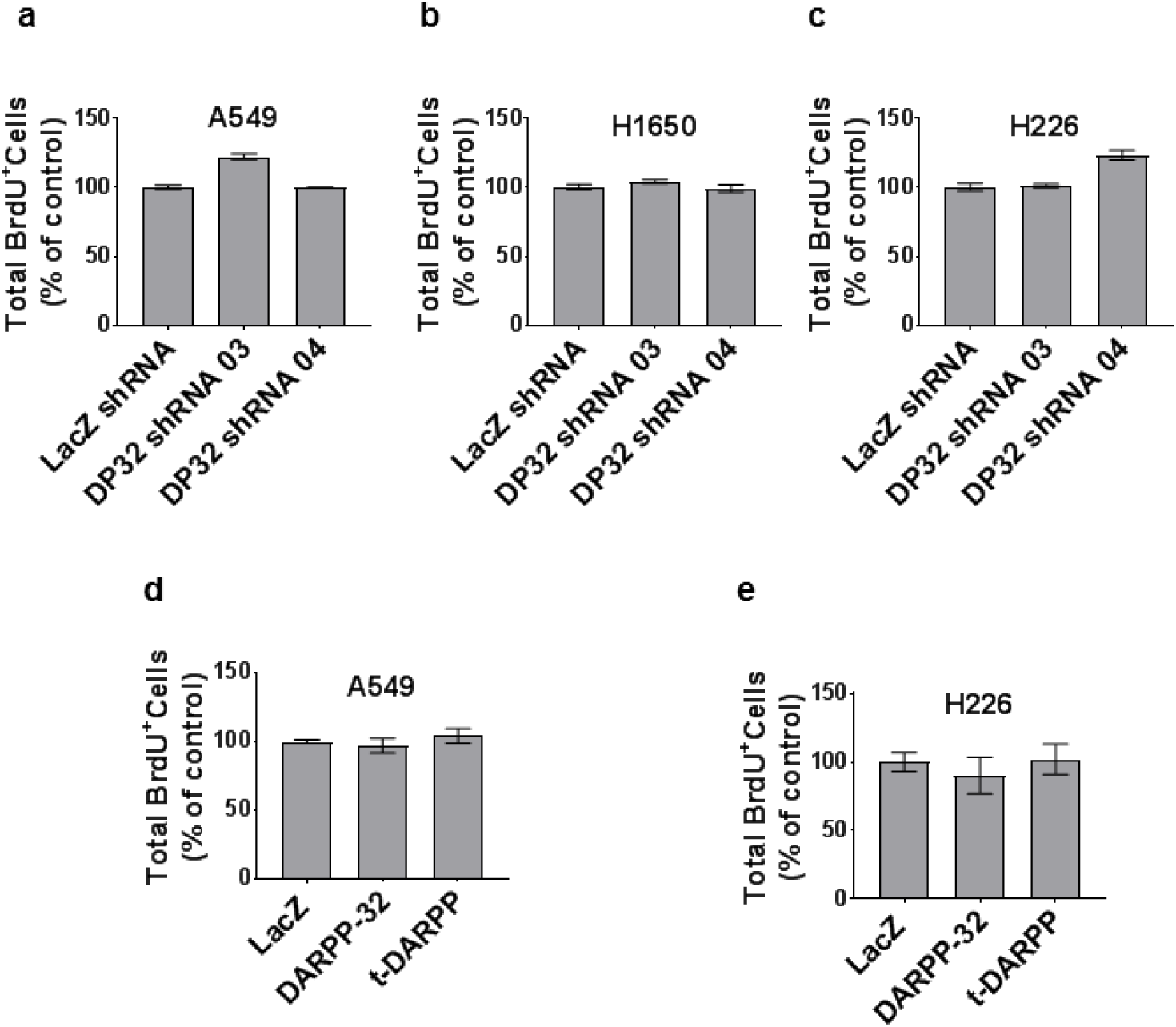
DARPP-32 does not regulate cell cycle progression. **a** A549, **b** H1650 and **c** H226 cells transduced with control or DARPP-32 shRNAs were seeded into 60-mm culture dishes for 16h. Flow cytometer-based BrdU cell proliferation assays were performed following incubation with anti-BrdU antibodies conjugated with APC. **d** A549 and **e** H226 cells were transduced with retrovirus containing control (LacZ), DARPP-32 or t-DARPP overexpressing clones. Flow cytometry-based BrdU cell proliferation assays were conducted and total BrdU+ cells were calculated. All bar graphs represent mean ± SE of at least 3 independent experiments.

**Supplementary Fig. 2:**
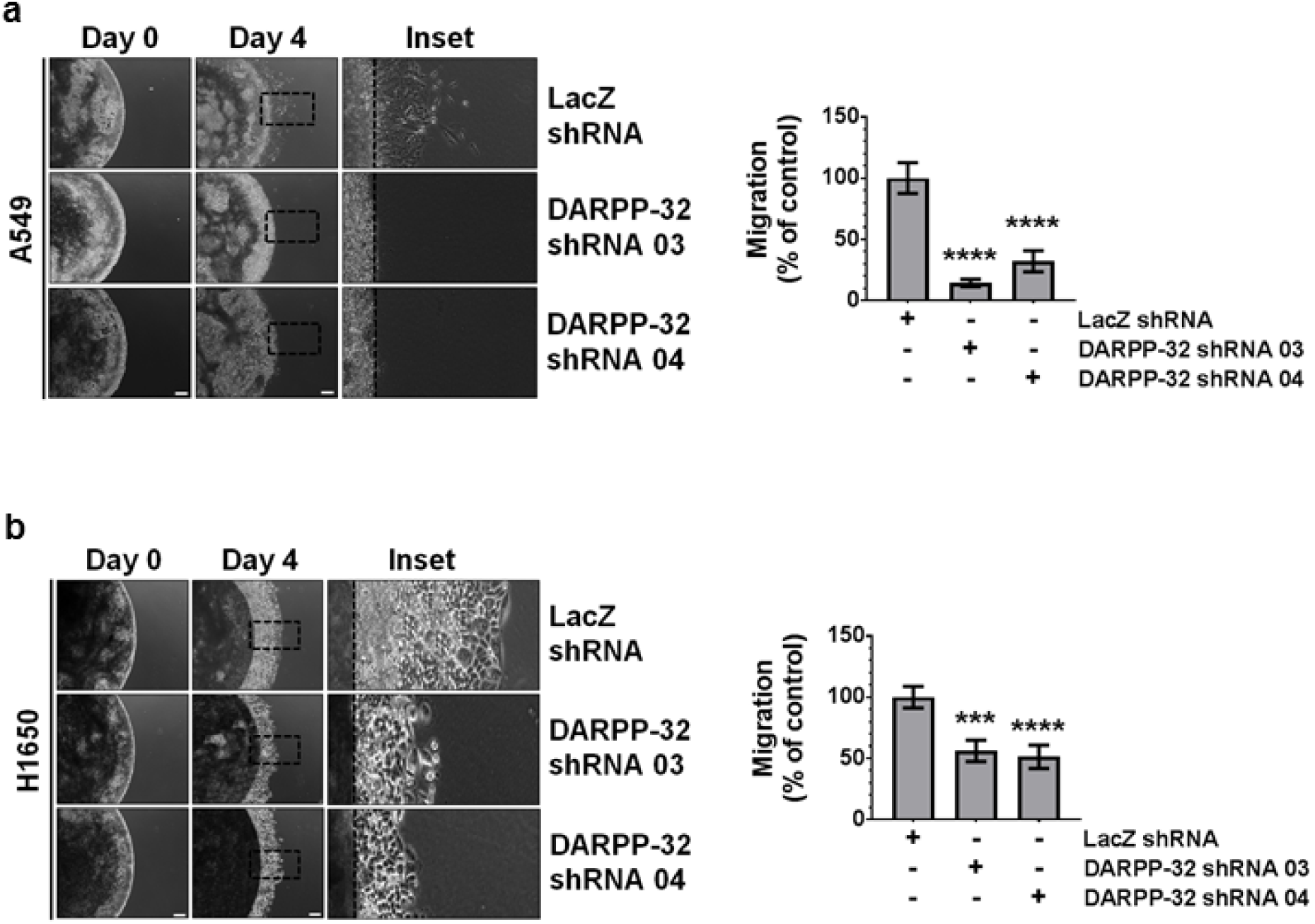
Knockdown of DARPP-32 reduces cell migration by spot assay. **a** A549 and **b** H1650 cells transduced with lentivirus encoding control or DARPP-32 shRNAs were mixed with Matrigel and spotted. Representative images depict one half of the spot at 0 and 4 days post-plating. Enlarged insets of day 4 images are depicted. The dashed line indicates the edge of the spot at day 0. Experiments were repeated at least three times. Scale bar, 200 μm. Distance travelled by the migratory cells were calculated using ImageJ software. Results represent mean ± SE. \*\*\**P*<0.001 and \*\*\*\**P*<0.0001, one-way ANOVA.

**Supplementary Fig. 3:**
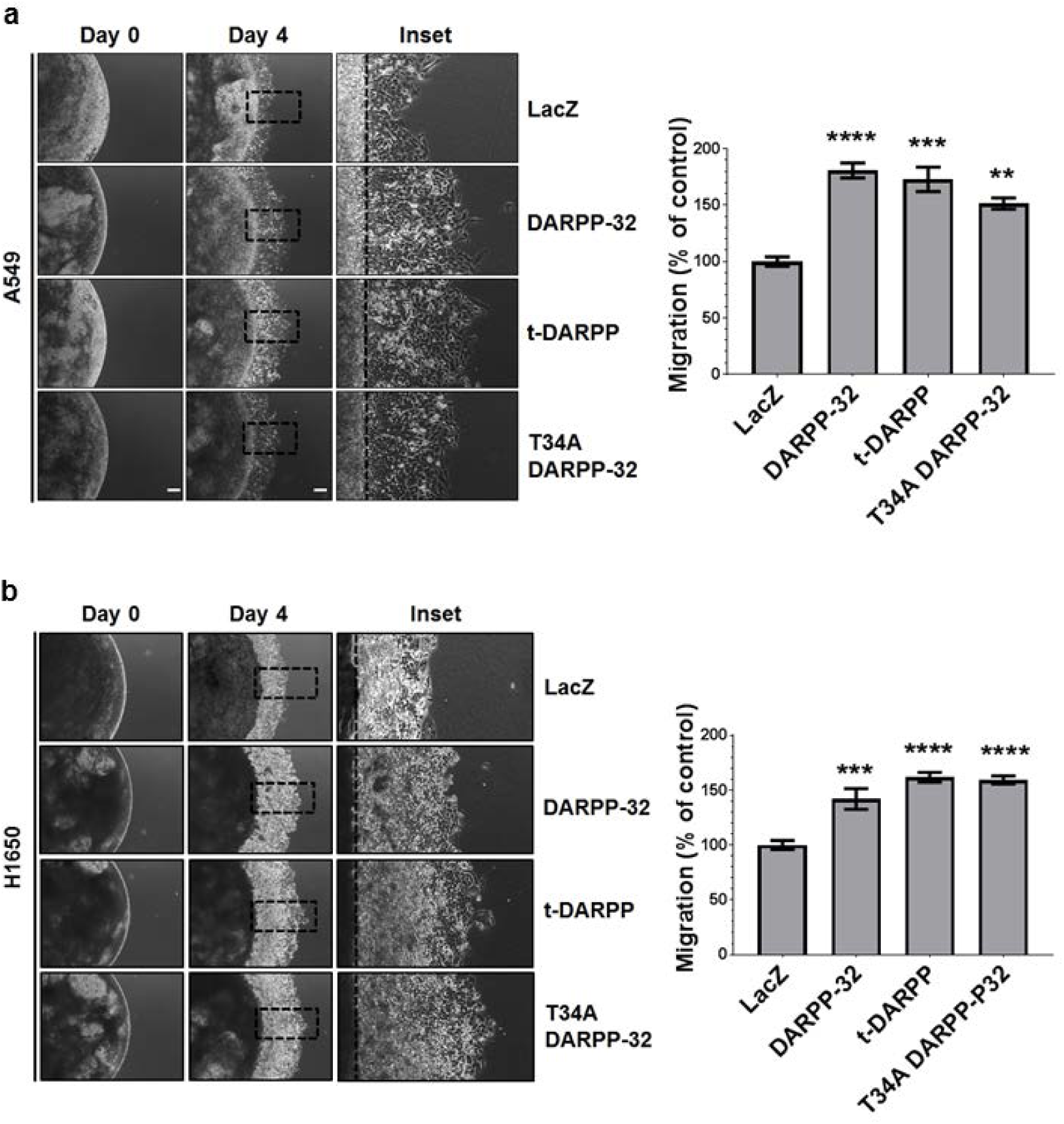
DARPP-32 increases cell migration by spot assay. **a** A549 and **b** H1650 cells transduced with retrovirus encoding control, DARPP-32, t-DARPP or T34A DARPP-32 clones were mixed with Matrigel and spotted. Representative images depict one half of the spot at 0 and 4 days post-plating. Enlarged insets of day 4 images are depicted. The dashed line indicates the edge of the spot at day 0. Experiments were repeated at least three times. Scale bar, 200 μm. Distance travelled by the migratory cells were calculated using ImageJ software. Results represent mean ± SE. \*\**P*<0.01, \*\*\**P*<0.001 and \*\*\*\**P*<0.0001, one-way ANOVA.

**Supplementary Fig. 4:**
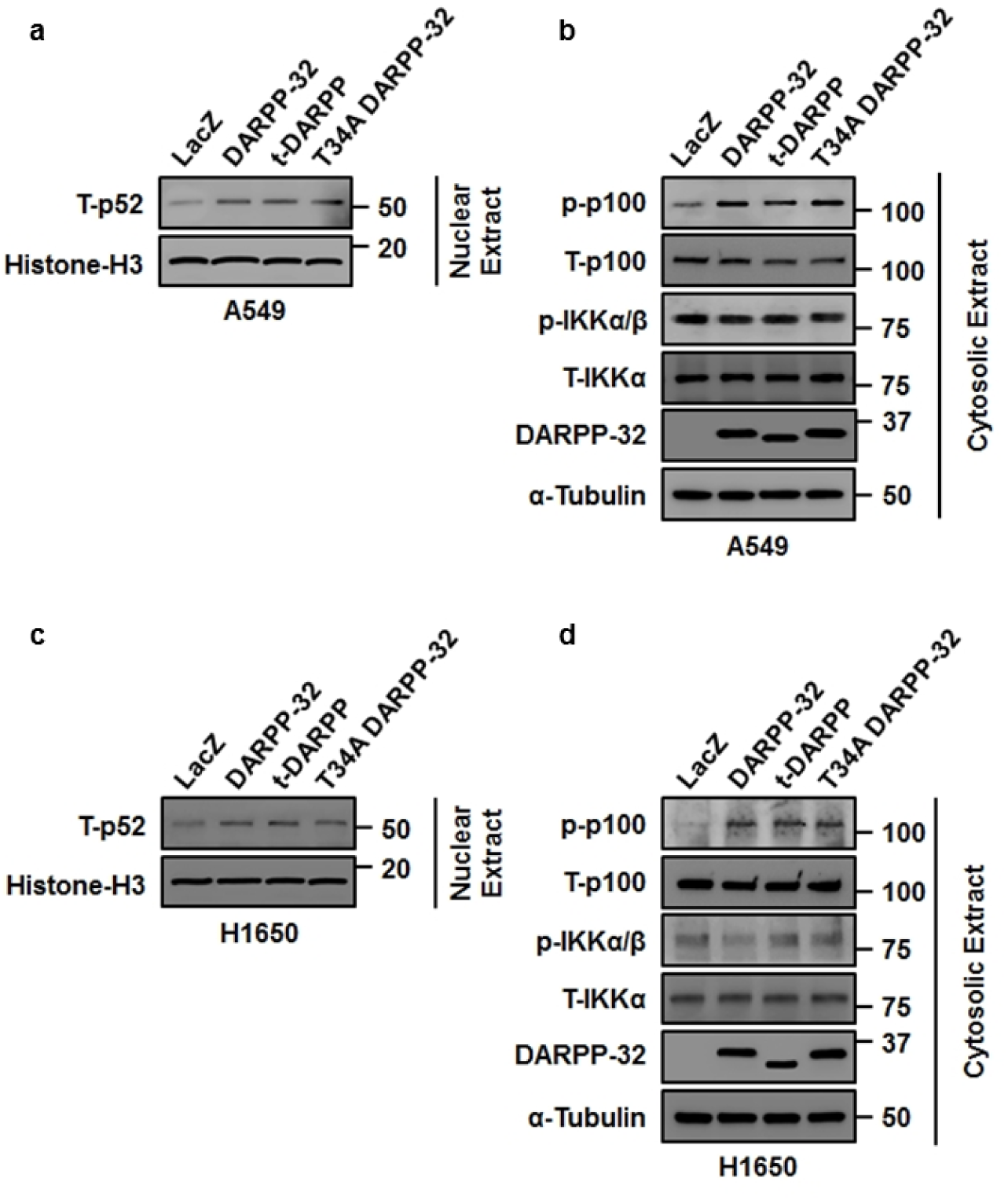
DARPP-32 positively regulates non-canonical NF-κB2 signaling. **a** Nuclear and **b** cytosolic fractions of A549 cells overexpressing LacZ control, DARPP-32, t-DARPP or T34A DARPP-32 clones were immunoblotted with antibodies against total p52 (T-p52), Histone H3 (loading control), phosphorylated p100 (p-p100), total p100 (T-p100), phosphorylated IKKα/β (p-IKKα/β), total IKKα (T-IKKα), DARPP-32 and a-tubulin (loading control). c Nuclear and d cytosolic fractions of H1650 cells overexpressing LacZ control, DARPP-32, t-DARPP or T34A DARPP-32 clones were subjected to western blotting using antibodies against total p52 (T-p52), Histone H3 (loading control), phosphorylated p100 (p-p100), total p100 (T-p100), phosphorylated IKKα/β (p-IKKα/β), total IKKα (T-IKKα), DARPP-32 and α-tubulin (loading control).

**Supplementary Fig. 5:**
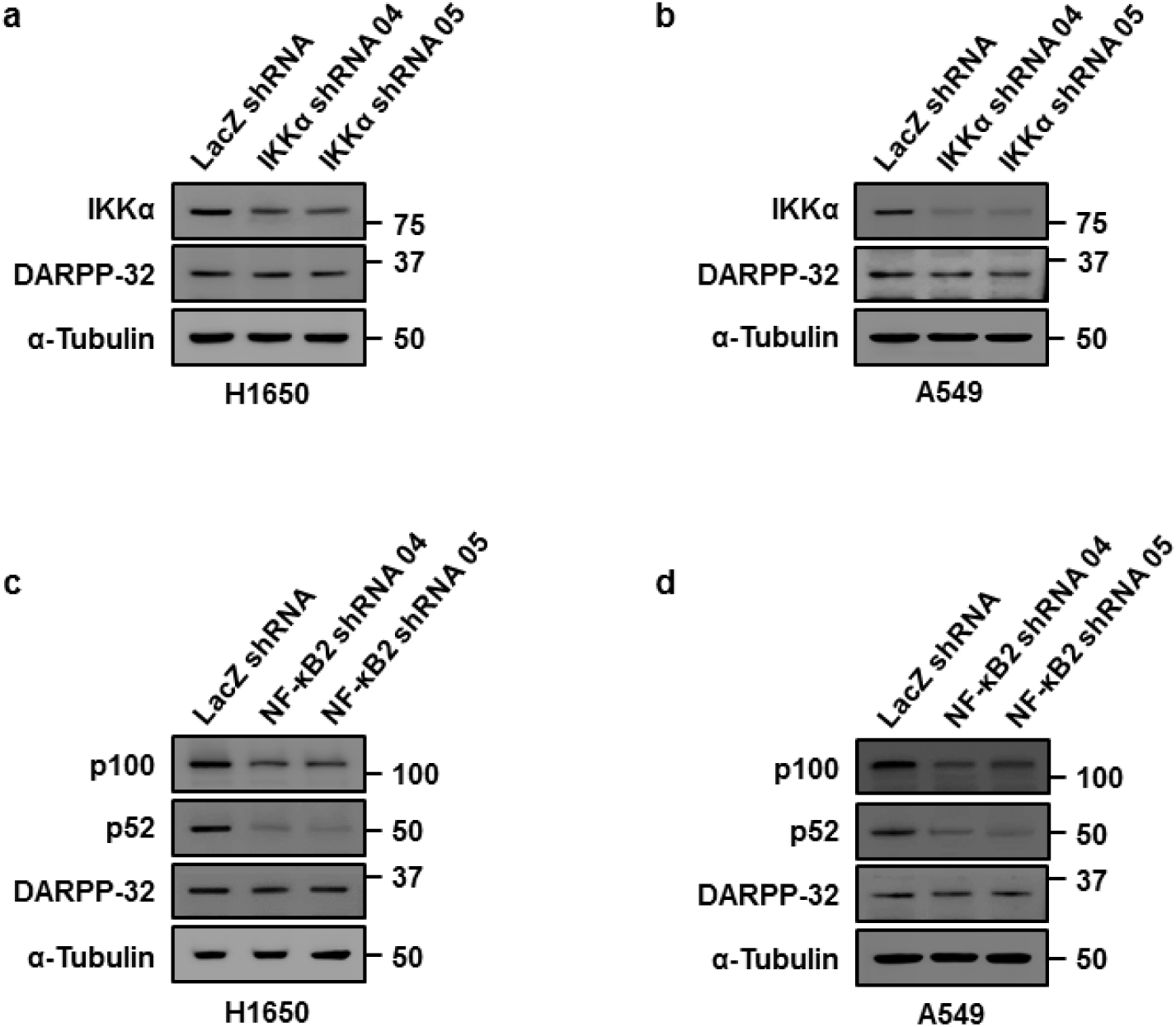
Abrogation of IKKα and NF-κB2 expression in NSCLC cells. **a** IKKα-depleted H1650 and **b** A549 cells were immunoblotted using antibodies against total IKKα, DARPP-32 and α-tubulin (loading control). c H1650 and d A549 cells were transduced with lentivirus encoding NF-κB2 shRNAs. Cell lysates were immunoblotted using antibodies against total NF-κB2 (detects both p100 and p52 subunits), DARPP-32 and α-tubulin (loading control).

**Supplementary Fig. 6:**
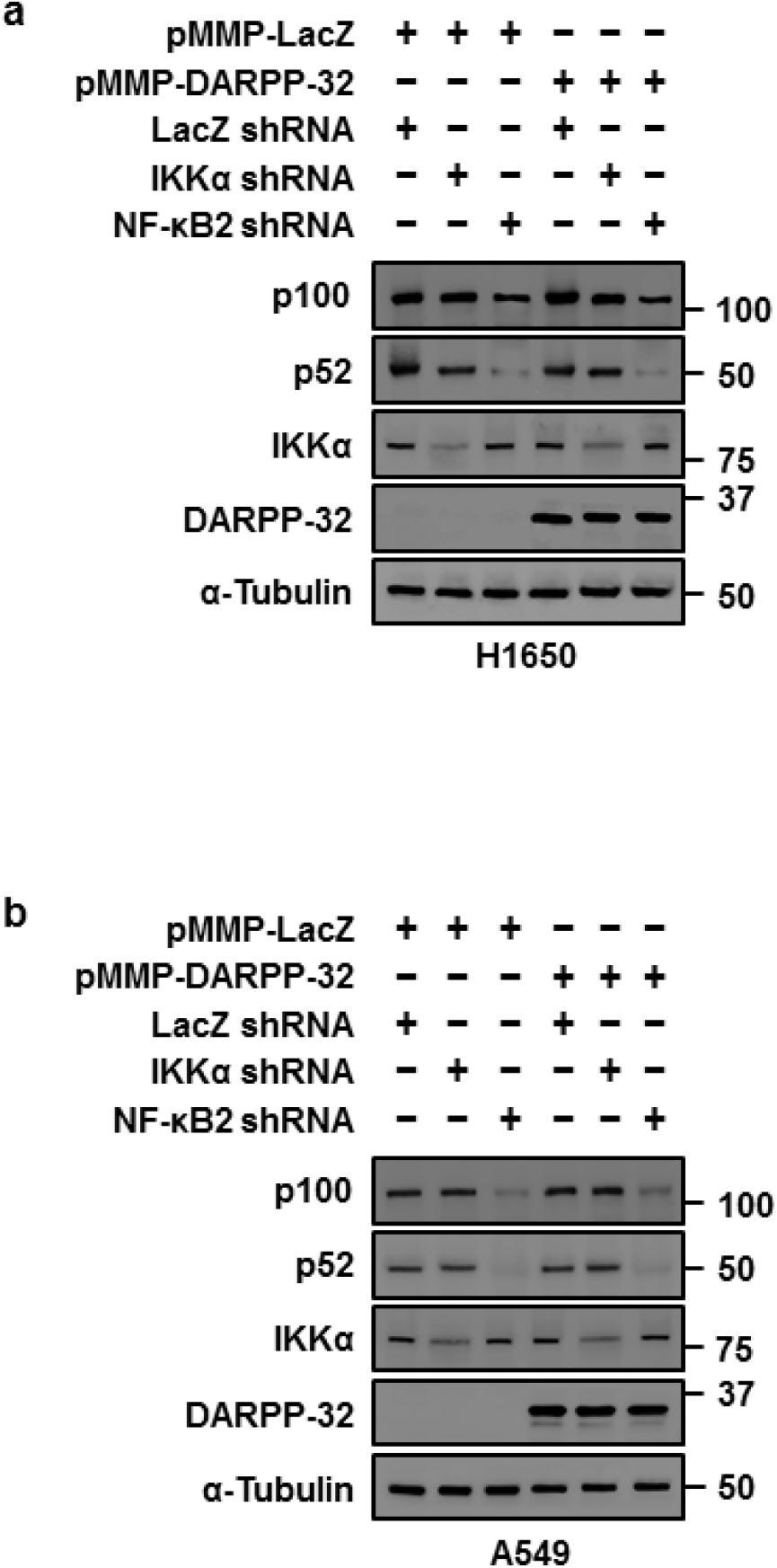
Overexpression of DARPP-32 in IKKα- and NF-κB2-depleted NSCLC cells. **a** H1650 and **b** A549 cells were transduced with lentivirus encoding IKKα or NF-κB2 shRNAs and then transfected with control (pMMP-LacZ) or DARPP-32 overexpressing plasmids (pMMP-DARPP-32) for 48 h. Cell lysates were harvested and immunoblotted using antibodies against total NF-κB2 (detects both p100 and p52 subunits), total IKKα, DARPP-32 and α-tubulin (loading control).

**Supplementary Fig. 7:**
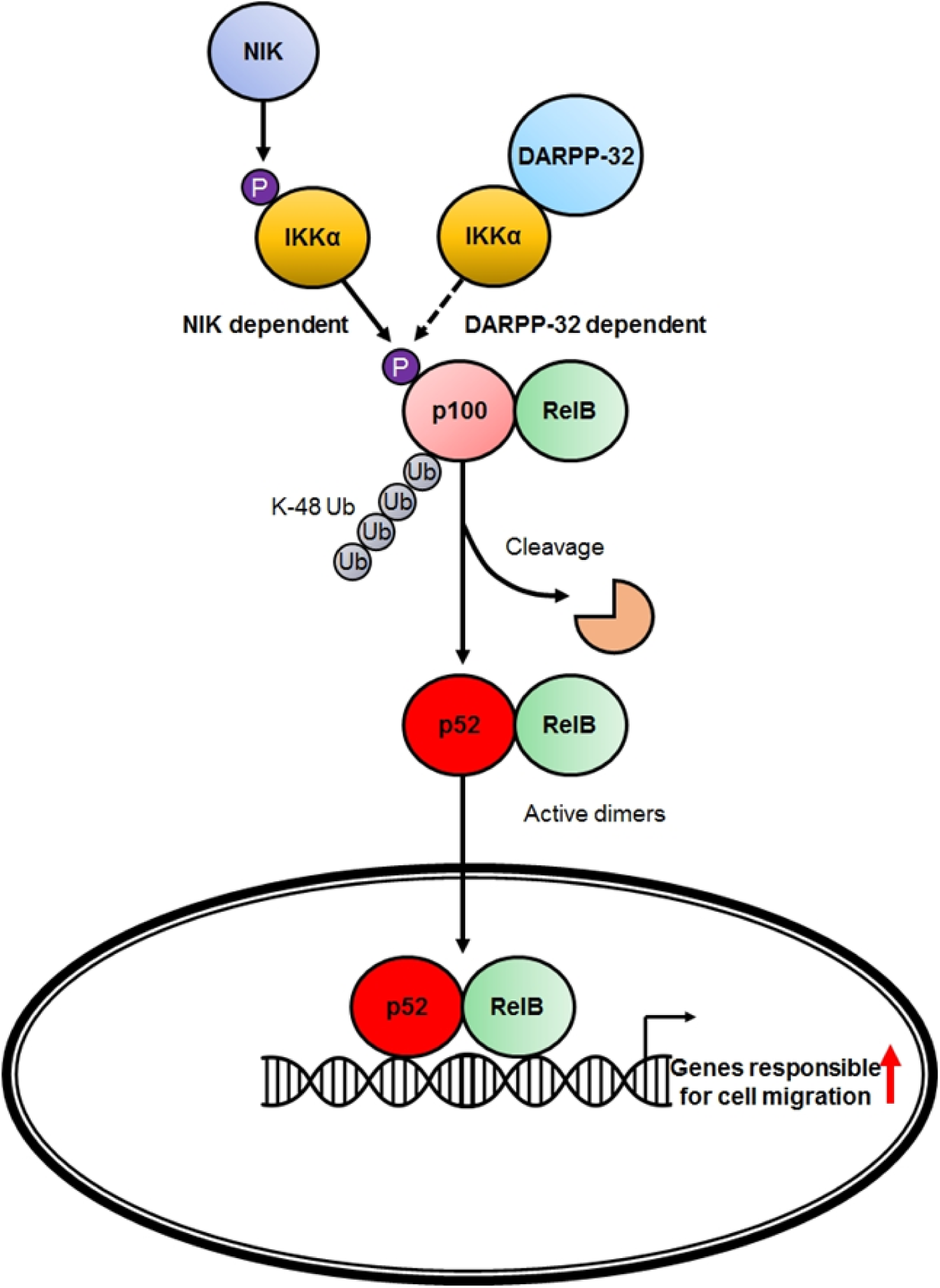
DARPP-32 promotes lung cancer cell migration through regulation of non-canonical NF-κB2 signaling. Upon stimulation, NF-κB inducing kinase (NIK) phosphorylates IKKα, which facilitates IKKα-mediated phosphorylation of p100 at S866/870. This phosphorylation event initiates subsequent ubiquitination at K48 and partial degradation of p100 by the proteasome to form p52. RelB-p52 heterodimers then translocate to the nucleus to regulate gene expression, including genes involved in cellular migration. Our data demonstrating a physical interaction between IKKα and DARPP-32 suggests IKKα-dependent phosphorylation of p100 may be directly or indirectly mediated by DARPP-32 through an NIK-independent mechanism (represented in dotted line) in human NSCLC cells.

**Supplementary Table 1:** Clinical characteristics of 62 lung adenocarcinoma patients corresponding to tissue specimens used to evaluate t-DARPP expression by immunohistochemistry

| CASE | GENDER | AGE | TUMOR | STAGE | DARPP-32 STAINING |  |  | DARPP-32 & t-DARPP STAINING |  |  |
| --- | --- | --- | --- | --- | --- | --- | --- | --- | --- | --- |
|  |  |  |  |  | Tumor Cell (%) | Intensity | IR Score | Tumor Cell (%) | Intensity | IR Score |
| 1 | MALE | 61 | T3 | IIIA | 90 | 3 | 270 | 100 | 3 | 300 |
| 2 | MALE | 64 | T3 | IIIA | 50 | 2 | 100 | 75 | 3 | 225 |
| 3 | MALE | 65 | T2 | IIIA | 0 | 0 | 0 | 0 | 0 | 0 |
| 4 | FEMALE | 66 | T1 | IIB | 0 | 0 | 0 | 1 | 2 | 2 |
| 5 | MALE | 72 | T1 | IIIA | 60 | 3 | 180 | 70 | 3 | 210 |
| 6 | FEMALE | 82 | T4 | IV | 90 | 3 | 270 | 95 | 3 | 285 |
| 7 | FEMALE | 80 | T3 | IIIB | 95 | 3 | 285 | 95 | 3 | 285 |
| 8 | FEMALE | 60 | T1 | IIIA | 100 | 3 | 300 | 100 | 3 | 300 |
| 9 | FEMALE | 72 | T1 | IIIA | 5 | 3 | 15 | 5 | 3 | 15 |
| 10 | MALE | 66 | T1 | IIIA | 0 | 0 | 0 | 2 | 3 | 6 |
| 11 | MALE | 71 | T2 | IIIA | 1 | 1 | 1 | 5 | 2 | 10 |
| 12 | FEMALE | 71 | T2 | IIIA | 100 | 3 | 300 | 100 | 3 | 300 |
| 13 | FEMALE | 72 | T3 | IIIA | 5 | 3 | 15 | 7 | 3 | 21 |
| 14 | MALE | 72 | T2 | IIIA | 95 | 3 | 285 | 95 | 3 | 285 |
| 15 | MALE | 64 | T1 | IIIA | 1 | 3 | 3 | 2 | 3 | 6 |
| 16 | MALE | 76 | T3 | IIIB | 100 | 3 | 300 | 100 | 3 | 300 |
| 17 | FEMALE | 51 | T3 | IIIA | 60 | 3 | 180 | 80 | 3 | 240 |
| 18 | MALE | 50 | T2 | IIIA | 100 | 3 | 300 | 100 | 3 | 300 |
| 19 | FEMALE | 52 | T1 | IIIA | 70 | 3 | 210 | 80 | 3 | 240 |
| 20 | FEMALE | 53 | T3 | IIIA | 5 | 3 | 15 | 10 | 3 | 30 |
| 21 | MALE | 76 | T3 | IIIB | 80 | 3 | 240 | 80 | 3 | 240 |
| 22 | FEMALE | 68 | T4 | IV | 80 | 3 | 240 | 90 | 3 | 270 |
| 23 | FEMALE | 47 | T3 | IIIA | 20 | 3 | 60 | 30 | 3 | 90 |
| 24 | FEMALE | 59 | T4 | IIIB | 0 | 0 | 0 | 0 | 0 | 0 |
| 25 | FEMALE | 36 | T2 | IIIA | 60 | 3 | 180 | 60 | 3 | 180 |
| 26 | FEMALE | 53 | T1 | IIIA | 30 | 3 | 90 | 60 | 3 | 180 |
| 27 | MALE | 66 | T3 | IIIA | 70 | 3 | 210 | 80 | 3 | 240 |
| 28 | MALE | 73 | T4 | IV | 70 | 3 | 210 | 85 | 3 | 255 |
|  |  |  |  |  | Tumor Cell (%) | Intensity | IR Score | Tumor Cell (%) | Intensity | IR Score |
| 29 | FEMALE | 56 | T1 | IIIA | 95 | 3 | 285 | 95 | 3 | 285 |
| 30 | FEMALE | 42 | T3 | IV | 0 | 0 | 0 | 0 | 0 | 0 |
| 31 | MALE | 71 | T2 | IIIA | 70 | 3 | 210 | 90 | 3 | 270 |
| 32 | MALE | 37 | T4 | IV | 100 | 3 | 300 | 100 | 3 | 300 |
| 33 | MALE | 57 | T1 | IIIA | 90 | 3 | 270 | 100 | 3 | 300 |
| 34 | MALE | 66 | T3 | IIIA | 20 | 3 | 60 | 40 | 3 | 120 |
| 35 | MALE | 54 | T4 | IIIB | 30 | 3 | 90 | 30 | 3 | 90 |
| 36 | MALE | 65 | T3 | IIIA | 60 | 3 | 180 | 100 | 3 | 300 |
| 37 | MALE | 55 | T2 | IIIA | 0 | 0 | 0 | 1 | 3 | 3 |
| 38 | MALE | 62 | T1 | IIIA | 70 | 3 | 210 | 85 | 3 | 255 |
| 39 | MALE | 49 | T2 | IIIA | 80 | 3 | 240 | 90 | 3 | 270 |
| 40 | FEMALE | 72 | T2 | IIIA | 80 | 3 | 240 | 100 | 3 | 300 |
| 41 | MALE | 65 | T3 | IIIA | 90 | 3 | 270 | 100 | 3 | 300 |
| 42 | MALE | 67 | T2 | IIIA | 0 | 0 | 0 | 1 | 3 | 3 |
| 43 | MALE | 69 | T1 | IIIA | 0 | 0 | 0 | 0 | 0 | 0 |
| 44 | MALE | 56 | T4 | IIIB | 0 | 0 | 0 | 1 | 3 | 3 |
| 45 | MALE | 67 | T1 | IIIA | 10 | 3 | 30 | 10 | 3 | 30 |
| 46 | MALE | 69 | T2 | IIIA | 1 | 3 | 3 | 1 | 3 | 3 |
| 47 | MALE | 76 | T4 | IV | 1 | 2 | 2 | 3 | 3 | 9 |
| 48 | MALE | 72 | T4 | IV | 100 | 3 | 300 | 100 | 3 | 300 |
| 49 | FEMALE | 60 | T1 | IIIA | 90 | 3 | 270 | 90 | 3 | 270 |
| 50 | FEMALE | 70 | T1 | IIIA | 10 | 3 | 30 | 60 | 3 | 180 |
| 51 | MALE | 60 | T3 | IIIA | 60 | 3 | 180 | 60 | 3 | 180 |
| 52 | FEMALE | 60 | T2 | IIIA | 10 | 3 | 30 | 10 | 3 | 30 |
| 53 | FEMALE | 73 | T2 | IIIA | 60 | 3 | 180 | 70 | 3 | 210 |
| 54 | MALE | 59 | T3 | IIIA | 0 | 0 | 0 | 0 | 0 | 0 |
| 55 | FEMALE | 63 | T3 | IIIA | 95 | 3 | 285 | 95 | 3 | 285 |
| 56 | MALE | 63 | T3 | IIIB | <1 | 3 | 3 | 1 | 3 | 3 |
| 57 | MALE | 36 | T3 | IIIA | 0 | 0 | 0 | 0 | 0 | 0 |
| 58 | FEMALE | 65 | T2 | IIIA | 0 | 0 | 0 | 1 | 3 | 3 |
| 59 | MALE | 55 | T2 | IIIA | 10 | 3 | 30 | 30 | 3 | 90 |
|  |  |  |  |  | Tumor Cell (%) | Intensity | IR Score | Tumor Cell (%) | Intensity | IR Score |
| 60 | MALE | 53 | T3 | IIIA | <1 | 3 | 3 | 1 | 3 | 3 |
| 61 | MALE | 65 | T1 | IIIA | 0 | 0 | 0 | 1 | 3 | 3 |
| 62 | MALE | 50 | T4 | IIIA | 30 | 3 | 90 | 60 | 3 | 180 |

## References

Alam, S. K., Yadav, V. K., Bajaj, S., Datta, A., Dutta, S. K., Bhattacharyya, M., Bhattacharya, S., Debnath, S., Roy, S., Boardman, L. A., Smyrk, T. C., Molina, J. R., Chakrabarti, S., Chowdhury, S., Mukhopadhyay, D., & Roychoudhury, S. (2016). DNA damage-induced ephrin-B2 reverse signaling promotes chemoresistance and drives EMT in colorectal carcinoma harboring mutant p53. Cell Death Differ, 23(4), 707–722. doi:10.1038/cdd.2015.133

Baldwin, A. S. (2001). Control of oncogenesis and cancer therapy resistance by the transcription factor NF-kappaB. J Clin Invest, 107(3), 241–246. doi: 10.1172/jci11991

Bang, D., Wilson, W., Ryan, M., Yeh, J. J., & Baldwin, A. S. (2013). GSK-3alpha promotes oncogenic KRAS function in pancreatic cancer via TAK1-TAB stabilization and regulation of noncanonical NF-kappaB. Cancer Discov, 3(6), 690–703. doi:10.1158/2159-8290.cd-12-0541

Beckler, A., Moskaluk, C. A., Zaika, A., Hampton, G. M., Powell, S. M., Frierson, H. F., Jr., & El-Rifai, W. (2003). Overexpression of the 32-kilodalton dopamine and cyclic adenosine 3’,5’-monophosphate-regulated phosphoprotein in common adenocarcinomas. Cancer, 98(7), 1547–1551. doi: 10.1002/cncr.11654

Beinke, S., & Ley, S. C. (2004). Functions of NF-kappaB1 and NF-kappaB2 in immune cell biology. Biochem J, 382(Pt 2), 393–409. doi:10.1042/bj20040544

Belkhiri, A., Dar, A. A., Peng, D. F., Razvi, M. H., Rinehart, C., Arteaga, C. L., & El-Rifai, W. (2008). Expression of t-DARPP mediates trastuzumab resistance in breast cancer cells. Clin Cancer Res, 14(14), 4564–4571. doi:10.1158/1078-0432.ccr-08-0121

Belkhiri, A., Dar, A. A., Zaika, A., Kelley, M., & El-Rifai, W. (2008). t-Darpp promotes cancer cell survival by up-regulation of Bcl2 through Akt-dependent mechanism. Cancer Res, 68(2), 395–403. doi: 10.1158/0008-5472.can-07-1580

Belkhiri, A., Zaika, A., Pidkovka, N., Knuutila, S., Moskaluk, C., & El-Rifai, W. (2005). Darpp-32: a novel antiapoptotic gene in upper gastrointestinal carcinomas. Cancer Res, 65(15), 6583–6592. doi:10.1158/0008-5472.CAN-05-1433

Belkhiri, A., Zhu, S., Chen, Z., Soutto, M., & El-Rifai, W. (2012). Resistance to TRAIL is mediated by DARPP-32 in gastric cancer. Clin Cancer Res, 18(14), 3889–3900. doi:10.1158/1078-0432.CCR-11-3182

Belkhiri, A., Zhu, S., & El-Rifai, W. (2016). DARPP-32: from neurotransmission to cancer. Oncotarget. doi:10.18632/oncotarget.7268

Bibb, J. A., Snyder, G. L., Nishi, A., Yan, Z., Meijer, L., Fienberg, A. A., Tsai, L. H., Kwon, Y. T., Girault, J. A., Czernik, A. J., Huganir, R. L., Hemmings, H. C., Jr., Nairn, A. C., & Greengard, P. (1999). Phosphorylation of DARPP-32 by Cdk5 modulates dopamine signalling in neurons. Nature, 402(6762), 669–671. doi:10.1038/45251

Caamano, J., & Hunter, C. A. (2002). NF-kappaB family of transcription factors: central regulators of innate and adaptive immune functions. Clin Microbiol Rev, 15(3), 414–429.

Cerami, E., Gao, J., Dogrusoz, U., Gross, B. E., Sumer, S. O., Aksoy, B. A., Jacobsen, A., Byrne, C. J., Heuer, M. L., Larsson, E., Antipin, Y., Reva, B., Goldberg, A. P., Sander, C., & Schultz, N. (2012). The cBio cancer genomics portal: an open platform for exploring multidimensional cancer genomics data. Cancer Discov, 2(5), 401–404. doi:10.1158/2159-8290.cd-12-0095

Cetin, K., Ettinger, D. S., Hei, Y., & O’Malley, C. D. (2011). Survival by histologic subtype in stage IV nonsmall cell lung cancer based on data from the Surveillance, Epidemiology and End Results Program. Clin Epidemiol, 3, 139–148. doi:10.2147/clep.s17191

Chen, Z., Zhu, S., Hong, J., Soutto, M., Peng, D., Belkhiri, A., Xu, Z., & El-Rifai, W. (2016). Gastric tumour-derived ANGPT2 regulation by DARPP-32 promotes angiogenesis. Gut, 65(6), 925–934. doi: 10.1136/gutjnl-2014-308416

Chen, Z. J., Parent, L., & Maniatis, T. (1996). Site-specific phosphorylation of IkappaBalpha by a novel ubiquitination-dependent protein kinase activity. Cell, 84(6), 853–862.

Cho, K., Cho, M. H., Seo, J. H., Peak, J., Kong, K. H., Yoon, S. Y., & Kim, D. H. (2015). Calpain-mediated cleavage of DARPP-32 in Alzheimer’s disease. Aging Cell, 14(5), 878–886. doi:10.1111/acel.12374

Christenson, J. L., Denny, E. C., & Kane, S. E. (2015). t-Darpp overexpression in HER2-positive breast cancer confers a survival advantage in lapatinib. Oncotarget, 6(32), 33134–33145. doi:10.18632/oncotarget.5311

Christenson, J. L., & Kane, S. E. (2014). Darpp-32 and t-Darpp are differentially expressed in normal and malignant mouse mammary tissue. Mol Cancer, 13, 192. doi: 10.1186/1476-4598-13-192

Cildir, G., Low, K. C., & Tergaonkar, V. (2016). Noncanonical NF-kappaB Signaling in Health and Disease. Trends Mol Med, 22(5), 414–429. doi:10.1016/j.molmed.2016.03.002

Dejardin, E., Droin, N. M., Delhase, M., Haas, E., Cao, Y., Makris, C., Li, Z. W., Karin, M., Ware, C. F., & Green, D. R. (2002). The lymphotoxin-beta receptor induces different patterns of gene expression via two NF-kappaB pathways. Immunity, 17(4), 525–535.

DiDonato, J. A., Hayakawa, M., Rothwarf, D. M., Zandi, E., & Karin, M. (1997). A cytokine-responsive IkappaB kinase that activates the transcription factor NF-kappaB. Nature, 388(6642), 548–554. doi:10.1038/41493

El-Rifai, W., Smith, M. F., Jr., Li, G., Beckler, A., Carl, V. S., Montgomery, E., Knuutila, S., Moskaluk, C. A., Frierson, H. F., Jr., & Powell, S. M. (2002). Gastric cancers overexpress DARPP-32 and a novel isoform, t-DARPP. Cancer Res, 62(14), 4061–4064.

Gao, J., Aksoy, B. A., Dogrusoz, U., Dresdner, G., Gross, B., Sumer, S. O., Sun, Y., Jacobsen, A., Sinha, R., Larsson, E., Cerami, E., Sander, C., & Schultz, N. (2013). Integrative analysis of complex cancer genomics and clinical profiles using the cBioPortal. Sci Signal, 6(269), pl 1. doi:10.1126/scisignal.2004088

Greengard, P. (2001). The neurobiology of slow synaptic transmission. Science, 294(5544), 1024–1030. doi:10.1126/science.294.5544.1024

Gu, L., Waliany, S., & Kane, S. E. (2009). Darpp-32 and its truncated variant t-Darpp have antagonistic effects on breast cancer cell growth and herceptin resistance. PLoS One, 4(7), e6220. doi: 10.1371/journal.pone.0006220

Hamel, S., Bouchard, A., Ferrario, C., Hassan, S., Aguilar-Mahecha, A., Buchanan, M., Quenneville, L., Miller, W., & Basik, M. (2010). Both t-Darpp and DARPP-32 can cause resistance to trastuzumab in breast cancer cells and are frequently expressed in primary breast cancers. Breast Cancer Res Treat, 120(1), 47–57. doi:10.1007/s10549-009-0364-7

Hansen, C., Greengard, P., Nairn, A. C., Andersson, T., & Vogel, W. F. (2006). Phosphorylation of DARPP-32 regulates breast cancer cell migration downstream of the receptor tyrosine kinase DDR1. Exp Cell Res, 312(20), 4011–4018. doi:10.1016/j.yexcr.2006.09.003

Hansen, C., Howlin, J., Tengholm, A., Dyachok, O., Vogel, W. F., Nairn, A. C., Greengard, P., & Andersson, T. (2009). Wnt-5a-induced phosphorylation of DARPP-32 inhibits breast cancer cell migration in a CREB-dependent manner. J Biol Chem, 284(40), 27533–27543. doi:10.1074/jbc.M109.048884

Hayden, M. S., & Ghosh, S. (2008). Shared principles in NF-kappaB signaling. Cell, 132(3), 344–362. doi:10.1016/j.cell.2008.01.020

Hemmings, H. C., Jr., Greengard, P., Tung, H. Y., & Cohen, P. (1984). DARPP-32, a dopamine-regulated neuronal phosphoprotein, is a potent inhibitor of protein phosphatase-1. Nature, 310(5977), 503–505.

Hoeppner, L. H., Wang, Y., Sharma, A., Javeed, N., Van Keulen, V. P., Wang, E., Yang, P., Roden, A. C., Peikert, T., Molina, J. R., & Mukhopadhyay, D. (2015). Dopamine D2 receptor agonists inhibit lung cancer progression by reducing angiogenesis and tumor infiltrating myeloid derived suppressor cells. Mol Oncol, 9(1), 270–281. doi:10.1016/j.molonc.2014.08.008

Hong, J., Katsha, A., Lu, P., Shyr, Y., Belkhiri, A., & El-Rifai, W. (2012). Regulation of ERBB2 receptor by t-DARPP mediates trastuzumab resistance in human esophageal adenocarcinoma. Cancer Res, 72(17), 4504–4514. doi: 10.1158/0008-5472.CAN-12-1119

Huang, H. B., Horiuchi, A., Watanabe, T., Shih, S. R., Tsay, H. J., Li, H. C., Greengard, P., & Nairn, A. C. (1999). Characterization of the inhibition of protein phosphatase-1 by DARPP-32 and inhibitor-2. J Biol Chem, 274(12), 7870–7878.

Kaur, H., Phillips-Mason, P. J., Burden-Gulley, S. M., Kerstetter-Fogle, A. E., Basilion, J. P., Sloan, A. E., & Brady-Kalnay, S. M. (2012). Cadherin-11, a marker of the mesenchymal phenotype, regulates glioblastoma cell migration and survival in vivo. Mol Cancer Res, 10(3), 293–304. doi:10.1158/1541-7786.mcr-11-0457

Kendellen, M. F., Bradford, J. W., Lawrence, C. L., Clark, K. S., & Baldwin, A. S. (2014). Canonical and non-canonical NF-kappaB signaling promotes breast cancer tumor-initiating cells. Oncogene, 33(10), 1297–1305. doi:10.1038/onc.2013.64

Kew, R. R., Penzo, M., Habiel, D. M., & Marcu, K. B. (2012). The IKKalpha-dependent NF-kappaB p52/RelB noncanonical pathway is essential to sustain a CXCL12 autocrine loop in cells migrating in response to HMGB1. J Immunol, 188(5), 2380–2386. doi:10.4049/jimmunol.1102454

Kopljar, M., Patrlj, L., Korolija-Marinic, D., Horzic, M., Cupurdija, K., & Bakota, B. (2015). High Expression of DARPP-32 in Colorectal Cancer Is Associated With Liver Metastases and Predicts Survival for Dukes A and B Patients: Results of a Pilot Study. Int Surg, 100(2), 213–220. doi:10.9738/intsurg-d-14-00022.1

Kunii, Y., Hyde, T. M., Ye, T., Li, C., Kolachana, B., Dickinson, D., Weinberger, D. R., Kleinman, J. E., & Lipska, B. K. (2014). Revisiting DARPP-32 in postmortem human brain: changes in schizophrenia and bipolar disorder and genetic associations with t-DARPP-32 expression. Mol Psychiatry, 19(2), 192–199. doi:10.1038/mp.2012.174

Liang, C. C., Park, A. Y., & Guan, J. L. (2007). In vitro scratch assay: a convenient and inexpensive method for analysis of cell migration in vitro. Nat Protoc, 2(2), 329–333. doi:10.1038/nprot.2007.30

Ling, J., Kang, Y., Zhao, R., Xia, Q., Lee, D. F., Chang, Z., Li, J., Peng, B., Fleming, J. B., Wang, H., Liu, J., Lemischka, I. R., Hung, M. C., & Chiao, P. J. (2012). KrasG12D-induced IKK2/beta/NF-kappaB activation by IL-1 alpha and p62 feedforward loops is required for development of pancreatic ductal adenocarcinoma. Cancer Cell, 21(1), 105–120. doi:10.1016/j.ccr.2011.12.006

Mirsadraee, S., Oswal, D., Alizadeh, Y., Caulo, A., & Van Beek, E. J. R. (2012). The 7th lung cancer TNM classification and staging system: Review of the changes and implications. World J Radiol, 4(4), 128–134. doi:10.4329/wjr.v4.i4.128

Molina, J. R., Yang, P., Cassivi, S. D., Schild, S. E., & Adjei, A. A. (2008). Non-small cell lung cancer: epidemiology, risk factors, treatment, and survivorship. Mayo Clin Proc, 83(5), 584–594. doi:10.4065/83.5.584

Rinkenbaugh, A. L., Cogswell, P. C., Calamini, B., Dunn, D. E., Persson, A. I., Weiss, W. A., Lo, D. C., & Baldwin, A. S. (2016). IKK/NF-kappaB signaling contributes to glioblastoma stem cell maintenance. Oncotarget, 7(43), 69173–69187. doi:10.18632/oncotarget.12507

Rotow, J., & Bivona, T. G. (2017). Understanding and targeting resistance mechanisms in NSCLC. Nat Rev Cancer, 17(11), 637–658. doi:10.1038/nrc.2017.84

Senftleben, U., Cao, Y., Xiao, G., Greten, F. R., Krahn, G., Bonizzi, G., Chen, Y., Hu, Y., Fong, A., Sun, S. C., & Karin, M. (2001). Activation by IKKalpha of a second, evolutionary conserved, NF-kappa B signaling pathway. Science, 293(5534), 1495–1499. doi:10.1126/science.1062677

Siegel, R. L., Miller, K. D., & Jemal, A. (2017). Cancer statistics, 2017. CA: A Cancer Journal for Clinicians, 67(1), 7–30. doi:10.3322/caac.21387

Torre, L. A., Siegel, R. L., & Jemal, A. (2016). Lung Cancer Statistics. Adv Exp Med Biol, 893, 1–19. doi: 10.1007/978-3-319-24223-1 1

Vangamudi, B., Peng, D. F., Cai, Q., El-Rifai, W., Zheng, W., & Belkhiri, A. (2010). t-DARPP regulates phosphatidylinositol-3-kinase-dependent cell growth in breast cancer. Mol Cancer, 9, 240. doi:10.1186/1476-4598-9-240

Vangamudi, B., Zhu, S., Soutto, M., Belkhiri, A., & El-Rifai, W. (2011). Regulation of beta-catenin by t-DARPP in upper gastrointestinal cancer cells. Mol Cancer, 10, 32. doi: 10.1186/1476-4598-10-32

Walaas, S. I., Nairn, A. C., & Greengard, P. (1983). Regional distribution of calcium- and cyclic adenosine 3’:5’-monophosphate-regulated protein phosphorylation systems in mammalian brain. I. Particulate systems. JNeurosci, 3(2), 291–301.

Wang, M. S., Pan, Y., Liu, N., Guo, C., Hong, L., & Fan, D. (2005). Overexpression of DARPP-32 in colorectal adenocarcinoma. Int J Clin Pract, 59(1), 58–61. doi:10.1111/j.1742-1241.2004.00305.x

Yang, J., Amiri, K. I., Burke, J. R., Schmid, J. A., & Richmond, A. (2006). BMS-345541 targets inhibitor of kappaB kinase and induces apoptosis in melanoma: involvement of nuclear factor kappaB and mitochondria pathways. Clin Cancer Res, 12(3 Pt 1), 950–960. doi: 10.1158/1078-0432.ccr-05-1220

Yeudall, W. A., Vaughan, C. A., Miyazaki, H., Ramamoorthy, M., Choi, M. Y., Chapman, C. G., Wang, H., Black, E., Bulysheva, A. A., Deb, S. P., Windle, B., & Deb, S. (2012). Gain-of-function mutant p53 upregulates CXC chemokines and enhances cell migration. Carcinogenesis, 33(2), 442–451. doi: 10.1093/carcin/bgr270

Zeng, H., Dvorak, H. F., & Mukhopadhyay, D. (2001). Vascular permeability factor (VPF)/vascular endothelial growth factor (VEGF) peceptor-1 down-modulates VPF/VEGF receptor-2-mediated endothelial cell proliferation, but not migration, through phosphatidylinositol 3-kinase-dependent pathways. J Biol Chem, 276(29), 26969–26979. doi:10.1074/jbc.M103213200

Zhu, S., Belkhiri, A., & El-Rifai, W. (2011). DARPP-32 increases interactions between epidermal growth factor receptor and ERBB3 to promote tumor resistance to gefitinib. Gastroenterology, 141(5), 1738–1748 e1731-1732. doi:10.1053/j.gastro.2011.06.070

Zhu, S., Chen, Z., Katsha, A., Hong, J., Belkhiri, A., & El-Rifai, W. (2016). Regulation of CD44E by DARPP-32-dependent activation of SRp20 splicing factor in gastric tumorigenesis. Oncogene, 35(14), 1847–1856. doi:10.1038/onc.2015.250

Zhu, S., Hong, J., Tripathi, M. K., Sehdev, V., Belkhiri, A., & El-Rifai, W. (2013). Regulation of CXCR4-mediated invasion by DARPP-32 in gastric cancer cells. Mol Cancer Res, 11(1), 86–94. doi:10.1158/1541-7786.mcr-12-0243-t

Zhu, S., Soutto, M., Chen, Z., Peng, D., Romero-Gallo, J., Krishna, U. S., Belkhiri, A., Washington, M. K., Peek, R., & El-Rifai, W. (2016). Helicobacter pylori-induced cell death is counteracted by NF-kappaB-mediated transcription of DARPP-32. Gut. doi:10.1136/gutjnl-2016-312141

